# Genome engineering by RNA-guided transposition for *Anabaena* PCC 7120

**DOI:** 10.1101/2022.09.18.508393

**Authors:** Sergio Arévalo, Daniel Pérez Rico, Dolores Abarca, Laura W. Dijkhuizen, Cristina Sarasa-Buisan, Peter Lindblad, Enrique Flores, Sandra Nierzwicki-Bauer, Henriette Schluepmann

**Affiliations:** Biology Department, Utrecht University, Utrecht, The Netherlands; Microbial chemistry, Department of Chemistry-Ångström Laboratory, Uppsala University, Uppsala, Sweden; Instituto de Bioquímica Vegetal y Fotosíntesis, CSIC and Universidad de Sevilla, Sevilla, Spain; Department of Biological Sciences, Rensselaer Polytechnic Institute, Troy, NY, USA; Department of Life Sciences, University of Alcalá, Alcalá de Henares, Spain

**Keywords:** *Anabaena*, CRISPR-associated transposon (CAST), genome engineering, RNA-guided transposition, minion sequencing, *de novo* genome assembly

## Abstract

In genome engineering, integration of incoming DNA has been dependent on enzymes produced by dividing cells which has been a bottle neck towards increasing DNA-insertion frequencies and accuracy. Recently, RNA-guided transposition with CRISPR-associated transposase (CAST) was reported as highly effective and specific in *Escherichia coli*. Here we developed Golden-Gate vectors to test CAST in filamentous cyanobacteria and show that it is effective in *Anabaena* sp. strain PCC 7120. The comparatively large plasmids containing the CAST and the engineered transposon were successfully transferred into *Anabaena* via conjugation using either suicide or replicative plasmids. Single guide(sg)-RNA encoding the leading, but not the reverse complement strand of the target were effective with the protospacer associated motif (PAM) sequence included in the sgRNA. In four out of six cases analyzed over two distinct target loci, the insertion site was exactly 63 bases after the PAM. CAST on a replicating plasmid was toxic which could be used to cure the plasmid. In all six cases analyzed, only the transposon cargo defined by the sequence ranging from left and right elements was inserted at the target loci, therefore, RNA-guided transposition resulted from cut and paste. No endogenous transposons were remobilized by exposure to CAST enzymes. This work is foundational for genome editing by RNA-guided transposition in filamentous cyanobacteria, whether in culture or in complex communities.

## Introduction

Cyanobacteria are of critical importance for the biogeochemical cycling of carbon and nitrogen and therefore substantially influence climate and primary production on earth (Sánchez-Baracaldo et al., 2022). Recent insights highlight the importance of symbioses in these cycles (Zehr and Capone, 2020). Indeed, cyanobacteria are capable of forming complex communities and they form symbioses with eukaryotic organisms including algae, plants, fungi, protozoa and invertebrate animals such as sponges and ascidians. This is in part because of their versatile secondary metabolism (Mutalipassi et al., 2021). Filamentous cyanobacteria from the order Nostocales typically produce O_2_-tight heterocysts that can fix dinitrogen in symbioses with plants from all land plant lineages (Rikkinen, 2017). Large sequencing datasets are growing in number expanding our understanding of those associations. However, understanding of biogeochemical interdependences at the molecular level within these associations has been hampered by the relative genetic intractability of the mostly polyploid cyanobacteria, whether in culture or in symbiotic communities.

Genetic alteration in filamentous cyanobacteria has been accomplished mostly in a few species from the *Nostoc/Anabaena* genus complex. DNA cargo transfer into these filamentous cells was achieved by natural competence, electroporation, *E. coli*-mediated conjugation and *Agrobacterium*-mediated transfer (Gutiérrez et al., 2021). Novel approaches for gene transfer have included the use of cell penetrating peptides. Stabilization of the incoming DNA was further achieved by methylation of the cargo in donor cells (Elhai et al., 1997), but also by engineering the DNA sequence such that the DNA may replicate and/or be used as a substrate for homologous recombination. This allowed for the integration of engineered DNA into target loci on either a plasmid or the chromosome. In all cases, integration of the cargo DNA was catalyzed by rate limiting enzymes from the target cell either by homology directed repair (HDR) or the predominant non-homologous end-joining repair. More recently, RNA-guided CRISPR-associated nucleases have been introduced in cyanobacteria to catalyze more efficiently changes at specific bases, or cause larger deletions at the target sites of the polyploid species of *Anabaena*, in spite of toxicity issues (Ungerer et al., 2016; Niu et al., 2018; Baldanta et al., 2022). However, the RNA-guided nucleases Cas9 or Cpf1 do not permit the tagging of those cells bearing the edit, whether by using a fluorescent- or selection-marker. Sequence- and species-directed gene-disruption with tagging is of particular importance to follow genome edited bacteria in complex mixtures.

Sequence-directed transposition of DNA allows gene-disruption whilst tagging; its efficiency has been demonstrated recently when catalyzing insertion of the cargo DNA with CRISPR-associated transposases. Two systems, the types 1-F and V-K, have been studied in *Escherichia coli* (Klompe et al., 2019; Strecker et al., 2019). The type I-F from *Vibrio cholerae* uses a multi-protein effector consisting of the type-characteristic Cas3 protein complexing with CASCADE (complex for antiviral defense)-proteins (Klompe et al., 2019). The type V-K uses the single effector protein Cas12k which was discovered in filamentous cyanobacteria, including *Scytonema hofmannii* (Strecker et al., 2019; Saito et al., 2021). In *E. coli*, the CAST increased the frequency of insertion of the donor DNA up to 80 % without selection. Importantly, it furthermore afforded its guided insertion because the effector protein binds small RNA that guide the transposition. The mechanism of transposition was not thoroughly investigated but both cut- and copy-paste have been reported (Strecker et al., 2019; Vo et al., 2021).

The CASCADE required more proteins to be expressed than the type V-K, did not transpose DNA cargo over 1 kbp in size at high efficiency, and reproducibly inserted it 46-55 bases after the PAM, yet in varying orientations (Klompe et al., 2019). In contrast, the type V-K from *Scytonema hofmanni* inserted cargo DNA up to 10 kbp long, 60-66 bp after the PAM in the orientation 5’ left-end (LE) to right end (RE) 3’at high frequency. The ends of the Tn7-derived type V-K cargo transposon consist of the 150 bp LE and the 90 bp RE, encoding 3 and 4 transposase binding sites respectively (Peters et al., 2001). The *S. hofmanni* V-K (*sh*CAST) system was plagued with significant off-target insertions (Strecker et al., 2019), and may be affected strongly by transcription at the target locus (Rodrigo González Linares et al., 2020). Transcripts likely titrate away the guide RNA in a way similar to DNA oligonucleotides or, possibly, the RNA-polymerase complex displaces the CAST complex (Bialk et al., 2015; Zhang et al., 2020).

Type V-K transposition was reconstructed *in vitro*; it required TnsB known to join the 3’ ends of Tn7 with target DNA, TnsC an ATP-dependent transposase activator known to form heptameric rings on DNA and TniQ known to recruit TnsC to the target DNA (Stellwagen and Craig, 1997; Peters and Craig, 2001; Park et al., 2021; Querques et al., 2021; Hoffmann et al., 2022). It also needed Cas12k which, like Cas9, is related to TnpB from the IS605 transposons (Faure et al., 2019). Unlike TnsA, Cas12k did not have the active site known to break the 5’ of the Tn7 ends; the Cas12k required two small RNAs: the constant transactivating RNA that formed a duplex with a part of the otherwise variant crRNA sequence. To specify new CAST targets with ease, the two small RNA binding to Cas12k have been expressed as a single-RNA (sgRNA) fusion and type two restriction enzyme cutting sites allow seamless cloning of the variable target sequence; the sgRNA design has been validated and optimized (Strecker et al., 2019).

Here we present the development of synthetic biology vectors with the elements from type V-K *sh*CAST comprised of the transposase proteins and the cargo transposon designed to study genome editing by way of RNA-guided and catalyzed insertion of transposon cargo in filamentous cyanobacteria. We tested RNA-guided transposition into highly expressed gene fusions in mutants of *Anabaena* sp. strain PCC 7120 (*Anabaena*), varying the target loci whilst keeping the sgRNA constant; we further varied the sgRNA sequence, PAM and strand. We characterized the insertions obtained in transconjugants by PCR, confocal microscopy and whole genome sequencing/*de novo* assembly to determine the mechanism of insertion and check for eventual off-target effects.

## Results

### CASTGATE elements to share for RNA-guided transposition in cyanobacteria

CAST elements were toxic when *E. coli* containing vectors encoding them were left on plates for over one week. To test toxicity and efficacy of combinations of CAST elements in cyanobacteria, they were transferred into the Golden Gate cloning vectors collectively called CASTGATE. This approach allows i) sharing of the individual elements in other synthetic biology studies (Table S1) and ii) assembling large vectors systematically with expression cassettes of CAST transposase proteins and the transposon cargo defined by the LE and RE (Figure 1).

**Figure 1.**
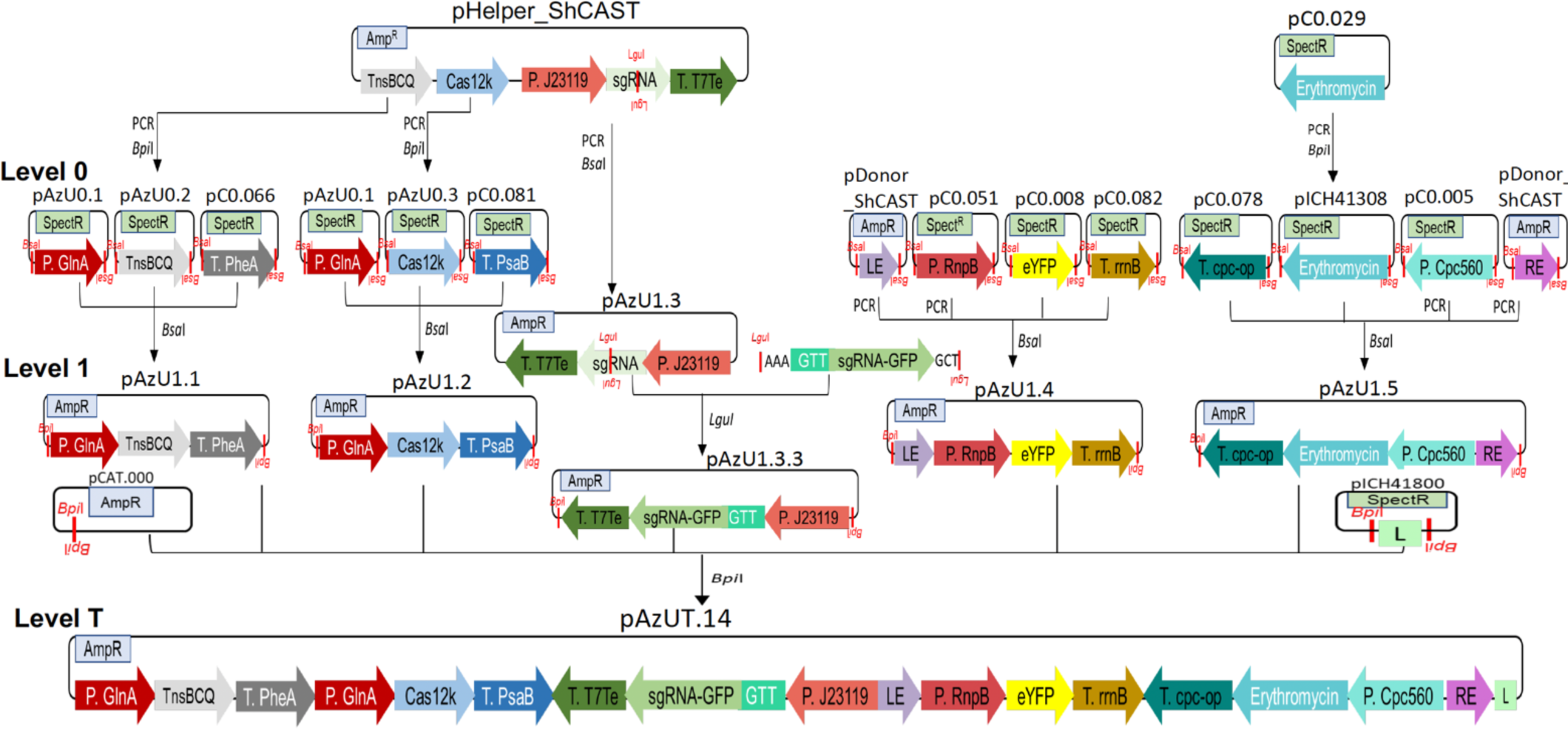
Cloning strategy to obtain the modules and vectors of the CASTGATE. The Level 0 plasmids with names beginning with pC0 and the donor for the Level T, pCAT.000, a replicative and conjugative vector were from the CyanoGate kit (Vasudevan R. at al., 2019). The level 0 pICH41308 and the Level 1 pICH41800 providing the linker (L) at position 6 in the T-level assemblies were from the MoClo kit (Weber et al., 2011). Firstly, sequences were domesticated and transferred in the Level 0 vectors: pAzU0.1 for P*glnA*, pAzU0.2 for the operon encoding TnsB, TnsC and TniQ (*tnsBCQ*), and pAzU0.3 for Cas12k which originated from pHelper_ShCAST (Strecker et al., 2019). Alternatively, sequences were PCR amplified for direct insertion into Level 1 vectors, as in the case of the cargo transposon LE and RE which originated from pDonor_ShCAST (Strecker et al., 2019), and the expression cassette of the sgRNA-scaffold from pHelper_ShCAST that allows to ligate target-specific sequences in LguI restriction sites. Secondly, expression cassettes were assembled in Level 1 for the specific positions (1.1 to 1.5) of the Level T assemblies: pAzU1.1, low nitrogen-inducible expression of the *tnsBCQ*; pAzU1.2, low nitrogen-inducible expression of Cas12k; pAzU1.3, the sgRNA-scaffold, which, when the GFP-specific target sequence (AAAGTT-GFPgRNA-GCT*) was ligated in the LguI site, yielded pAz1.3.3; pAzU1.4, with at the 5’ the LE, then the cassette for constitutive expression of eYFP; pAzU1.5, with at the 5’ the cassette for constitutive erythromycin resistance followed at the 3’ with the RE. Thirdly, the level 1 plasmids were combined to generate the Level T pAzUT.14, encoding the CAST machinery followed by the cargo DNA flanked by the LE and RE. CAST component Level 1 plasmids were replaced with linkers during the final Level T plasmid assemblies to test their individual toxicity and efficacy. * three different target sequences within the *gfp* sequence were tested as described in Figure S2.

Level 0 vectors generated for sharing contained the domesticated operon *tnsB, tnsC* and *tniQ,* the *cas12k*, and the LE and RE (Figure 1, Level 0). They further contained the *glnA* promoter from *Anabaena* (P*_glnA_*; Valladares et al., 2004) (Figure 1, *glnA*) used to increase expression when the cyanobacteria were grown without nitrogen in the culture medium (Figure 1). The level 1 vector for sgRNA expression was designed to transcribe away from the LE; it contained the sgRNA scaffold with LguI restriction sites allowing for the insertion of annealed primers specifying the sequence targeted by the sgRNA (Figure 1, level 1 position 3). The level T plasmids were assembled on backbones of conjugative and replicative vectors, non-conjugative replicative vectors and, conjugative but not replicative (suicide) vectors (Figure 1, level T).

The CASTGATE vectors listed in Table S1 proved stable in *E. coli*. We thus attempted transfer of the conjugative replicative vectors listed in Table S2 into *Anabaena* by triparental conjugation.

### RNA-guided transposition efficiently targets expressed genes irrespective of locus position in the *Anabaena* chromosome

Reference vectors containing only cargo DNA with expression cassettes for cytosolic eYFP (YFP) and erythromycin or spectinomycin/streptomycin-resistance (Table S1, pAzUT.3 and .4) yielded clones in all the *Anabaena* strains tested: the wild-type PCC7120, the derived strains containing the GFP fused to the ammonium transporter protein Amt1 (*amt1::gfp*, CSVT15; (Merino-Puerto et al., 2010) or the septal-protein SepJ (*sepJ::gfp*, CSAM137; (Flores et al., 2007).

On the nitrogen-rich BG11 medium, the following vectors yielded transconjugants: vectors containing all of the CAST and the cargo DNA but inactive sgRNA, those containing the sgRNA in isolation, or those with all the CAST elements and active sgRNA targeting the GFP sequence. Expression of the CAST components, therefore, was well enough repressed behind the P*_glnA_* on nitrogen rich medium, or the inherent toxicity of the CAST proteins was low enough for the transconjugant plasmids to be maintained in *Anabaena*.

On BG11_0_ medium without nitrogen, strain CSAM137 survived the short periods of induction for P*_glnA_-*dependent CAST-protein expression but not long periods. In contrast, strain CSVT15 grew on BG11_0_ which allowed to test toxicity of CAST proteins when P*_glnA_* strongly drives their expression (Figure S1). On BG11_0,_ Cas12k caused growth inhibition when expressed in isolation (Figure S1 28 d, pAzUT7) but when expressed with all the CAST components in transconjugants with pAzUT9 did not affect growth.

Homogenous clones cured of the donor plasmids were obtained at high frequency after sonication and growth on BG11 medium under erythromycin selection when using pAzUT14 in either the CSVT15 or CSAM137 strains. pAzUT14 expressed the sgRNA1 targeting the sense strand of the GFP 163 bp into the 563 bp protein (Figure S2).

Genotyping of the clones obtained at the *amt1::gfp* locus of CSVT15 was caried out by PCR amplification spanning either ends of the transposon (Figure 2A, PCR1, PCR2), the region targeted by the sgRNA (PCR3) and the pAzUT14 backbone (PCR4). In all three clones tested, the expected size fragments were amplified at either ends and the total size of the inserted transposons was consistent with a single 5’LE to RE3’ transposon insertion (Figure 2B, PCR1,2,3). The backbone of the donor vector could not be detected confirming that the clones were cured of it (Figure 2B, PCR4). Genotyping the clones obtained at the *sepJ::gfp* locus after curing pAzUT14 from transconjugants of CSAM137 gave a similar result (Figure 3A): amplicons spanning either ends (Figure 3B, PCR1, PCR2), amplicon spanning the entire length of the insertion (Figure 3B, PCR3) and absence of the backbone (Figure 3B, PCR4). Therefore, PCR-genotyping demonstrated that CAST was able to accurately guide the cargo transposon insertion into highly expressed recombinant GFP, independently of the *Anabaena* locus chosen.

**Figure 2.**
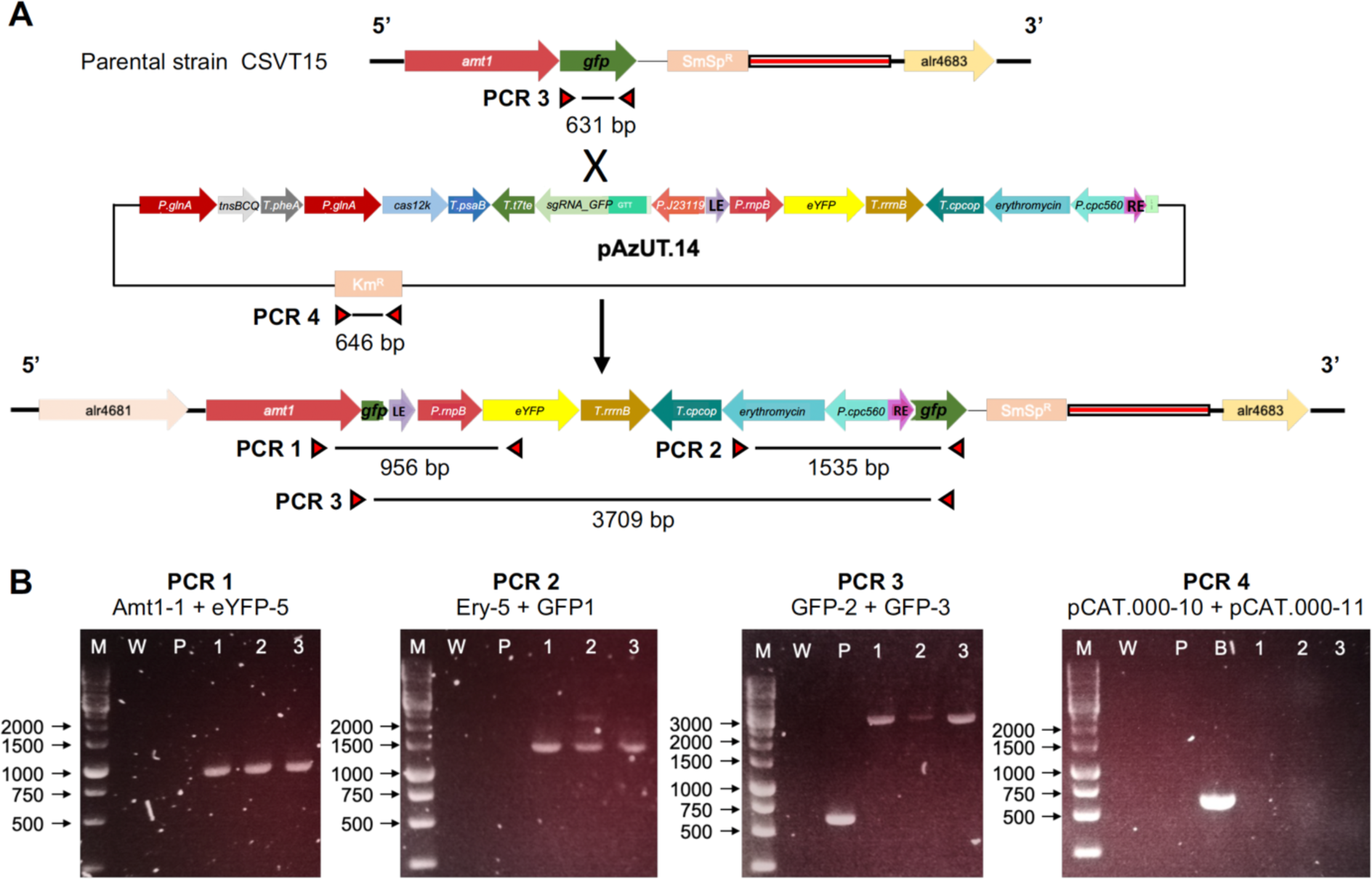
Scheme of the RNA-guided transposon insertion in the *gfp* of the *amt1::gfp* locus of the parental strain CSVT15, and its detection by PCR. **A,** Scheme of the *amt1::gfp* locus from strain CSVT15 that, after conjugation and selection on erythromycin, also contained pAzUT.14. pAzUT.14 encoded the RE- and LE-flanked transposon cargo, which would be mobilized by the separately encoded CAST enzymes and sgRNA targeting the *gfp*. When mobilized with complete resolution of the transposase complex, the cargo transposon inserts inside the *gfp* of the *amt1:gfp* fusion. **B,** PCR detection of cargo insertion and full segregation in the different exconjugant clones. PCR 1 detects the eYFP integration; PCR 2 detects erythromycin-resistance gene integration; PCR 3 detects *gfp* and thus segregation of transposed genotype; PCR 4 detects whether the exconjugant had been cured of pAzUT.14. The letters and numbers correspond with: (M) 1-kb DNA ladder; (W) *Anabaena* sp. PCC 7120; (P) CSVT15; (B) pAzUT.14; (1) AzUU1 clone 1; (2) AzUU1 clone 4; and (3) AzUU1 clone 8. *Anabaena* sp. PCC 7120 was used as negative control in all PCR tests. In the PCR 3, the CSVT15 strain was used as positive control.

**Figure 3.**
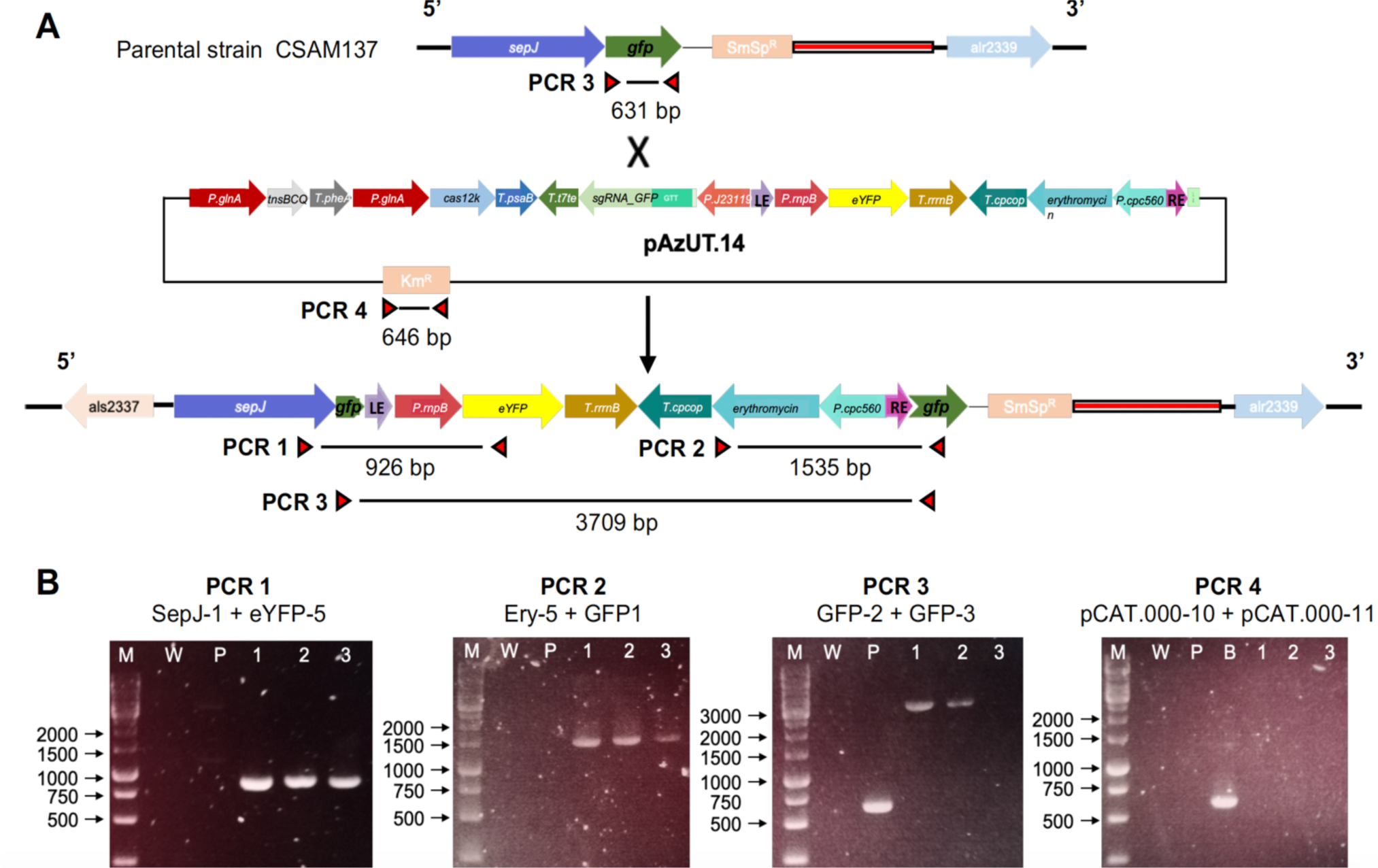
Scheme of the RNA-guided transposon insertion in the *gfp* of the *sepJ::gfp* locus of the parental strain CSAM137, and its detection a by PCR. **A**, Scheme of the *sepJ::gfp* locus from strain CSAM137 that, after conjugation and selection on erythromycin, also contained pAzUT.14. pAzUT.14 encoded the RE- and LE-flanked transposon cargo, which would be mobilized by the CAST enzymes and sgRNA targeting the *gfp,* also encoded in pAzUT.14. When mobilized with resolution of the transposase complex, the cargo transposon inserts inside the *gfp* of the *sepJ:gfp* fusion. **B**, PCR detection of cargo insertion and full segregation in the different exconjugants. PCR 1 detects the eYFP integration; PCR 2 detects erythromycin-resistance gene integration; PCR 3 detects *gfp* and thus segregation of transposed genotype; PCR 4 detects whether the exconjugant had been cured of pAzUT.14. The letters and numbers correspond with: (M) 1-kb DNA ladder; (W) *Anabaena* sp. PCC 7120; (P) CSAM137; (B) pAzUT.14; (1) AzUU2 clone 3; (2) AzUU2 clone 8; and (3) AzUU2 clone 7. *Anabaena* sp. PCC 7120 was used as negative control in all PCR tests. In the PCR 3, the CSAM137 strain was used as positive control.

Using the vector pAzUT18 which encodes an sgRNA targeting the sense strand at the wild-type locus *alr3727*, all transconjugant colonies tested by PCR, as described in Figure S3A, had traces of the cargo transposon insertion (Figure S3B) unlike the parental wild-type strain (Figure 3B, WT); which confirmed efficacy of the CAST on a wild-type gene.

### Integration of the cargo DNA inactivated the targeted GFP-fusions

Confocal viewing was not used for screening because it was not able to distinguish whether the fluorescence detected from the expression of YFP was encoded in the DNA cargo transposon on the donor plasmid or after insertion in a chromosomal locus. Nevertheless, in strains cured of the plasmids, it verified absence of the GFP-fusion protein (Figure 4 GFP) and strong YFP expression in the transconjugants obtained. YFP was seen as yellow fluorescence in the cytosol of all cells of the clones UU1 and UU2 (Figure 4 YFP) compared to the respective parents CSVT15 and CSAM137 which expressed the GFP fusions. This confirmed that the insertion of RNA-guided transposon cargo into the chromosome loci may be used to tag *Anabaena,* which is known to be polyploid (average number of chromosome 8.2, Hu et al., 2007).

**Figure 4.**
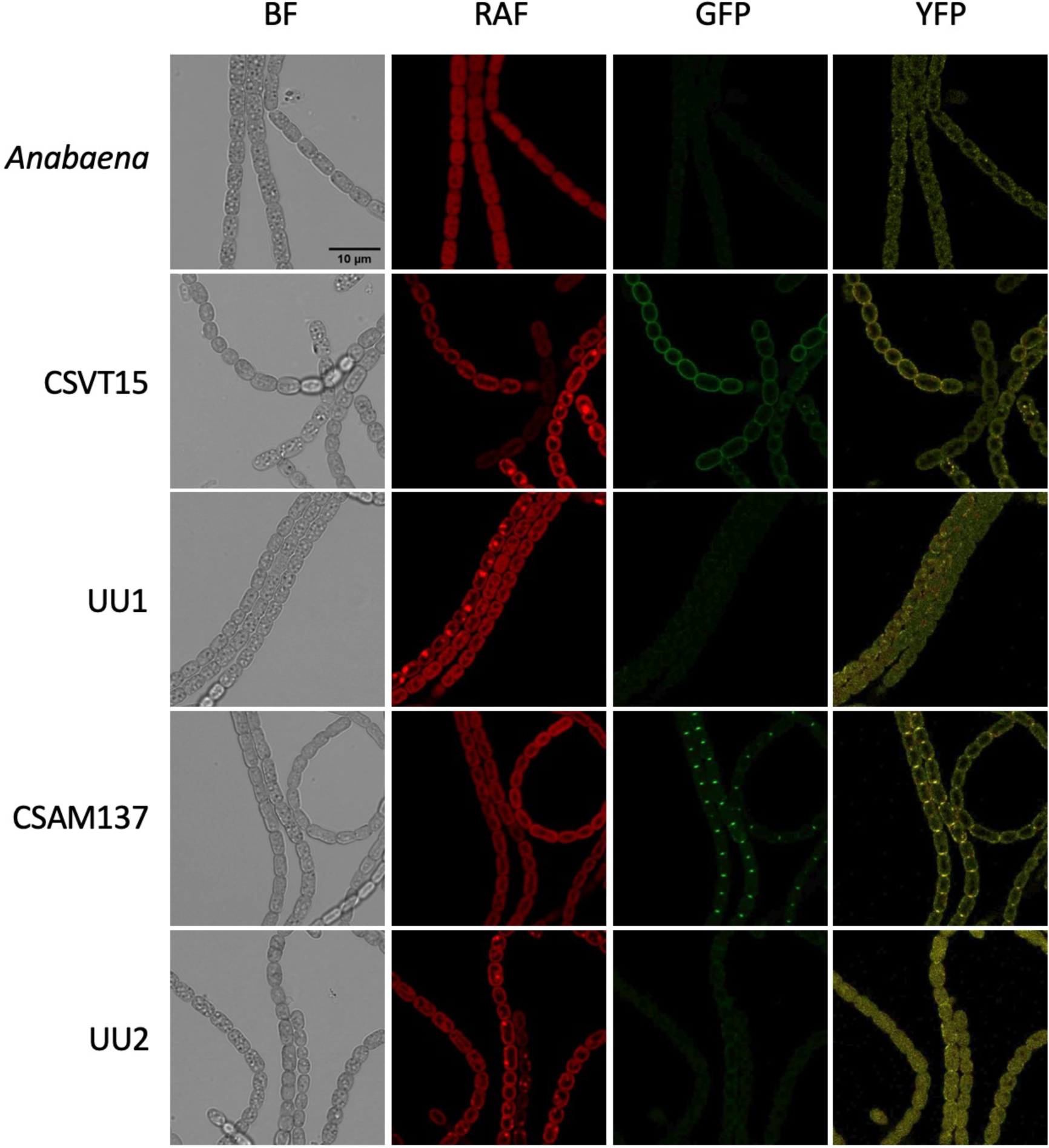
Fluorescence detection of the GFP-protein fusions and the transposon-encoded YFP in *Anabaena* strains. The strains UU1 and UU2 were fully segregated for the transposon insertion and cured of the pAzUT.14. Brightness and contrast settings were equal for all image detection types: the Bright Field (BF) and the fluorescence settings to detect Red Auto Fluorescence (RAF), Green Fluorescent Protein (GFP), and Yellow Fluorescent Protein (YFP). *Anabaena*, wild-type strain; CSVT15, the parental strain with the *amt1::gfp* locus; UU1, a CSVT15-derived exconjugant as shown in Fig. 2; CSAM137, the parental strain with the *sepJ::gfp* locus; UU2, a CSAM137 derived exconjugant as shown in Fig. 3. Scale bar, 10 μm. The images are representative of at least 20 micrographs for each strain gathered from two or three independent microscopy analyses.

We conclude therefore that the RNA-guided transposition in *Anabaena* may serve to generate tagged inactivation at expressed loci specified by the sgRNA.

### The sgRNA targeting the sense strand of expressed GFP and containing the GTT PAM was effective

We next explored how the sgRNA sequences affect the specificity and efficacy of RNA-guided transposition (Figure S2). The sgRNA from pAzUT14 efficiently targeted the sense-strand of the expressed *gfp* fusions and contained in its 5 prime the PAM GTT 162 bases into the *gfp* sequence (Figure 2, 3). Including the PAM in the sgRNA sequence for strong binding of the sgRNA to Cas12k was attempted because the sgRNA locus on the donor plasmid is protected by the proximity of the LE of the transposon from DNA cargo insertions and thus inactivation. Insertions of the cargo transposon were observed when sgRNA was used from pAzUT10 that contained the PAM GGTT and targeted the sense strand of *gfp*, 271 bp into the coding sequence of *gfp*, but these were recovered at much lower frequency (Figure S2, and data not shown). No insertions were observed for pAzUT12 where the sgRNA contained the same PAM GGTT and targeted the antisense strand of *gfp* 563 bp into the *gfp* sequence (Figure S2, and data not shown).

### The cargo transposon inserted mostly 63 b after the PAM and led to 2-5 base duplications at the insertion sites

PCR-results suggested that, in all cases, the cargo transposon was inserted in the 5’ LE to RE 3’orientation, but PCR could not distinguish whether the insertions had been a result of cut- or copy-paste mechanisms. For accurate sequence information on the integration loci, DNA was extracted from the six independently obtained clones exhibiting RNA-guided transposition (Figure 3 and 4), their respective parental strains, and from the wild-type *Anabaena*. The DNA was then sequenced using Minion long-reads with a minimum 50 x coverage which allowed assemblies to be resolved into a single full-length chromosome with high-accuracy sequence information.

The sequences immediately adjacent to the cargo transposon were extracted and aligned for comparison (Figure 5). In four of the six genomes analyzed, the LE inserted exactly 63 bases behind the PAM. When comparing sequences at the LE with RE, duplications of the bases at the vicinity of the insertion due to the resolution of transposase complex were two- to four bases long, and in five out of six cases they were exactly five bases (Figure 5, duplications highlighted in purple). RNA-guided cargo transposon insertions were therefore very reproducible.

**Figure 5.**
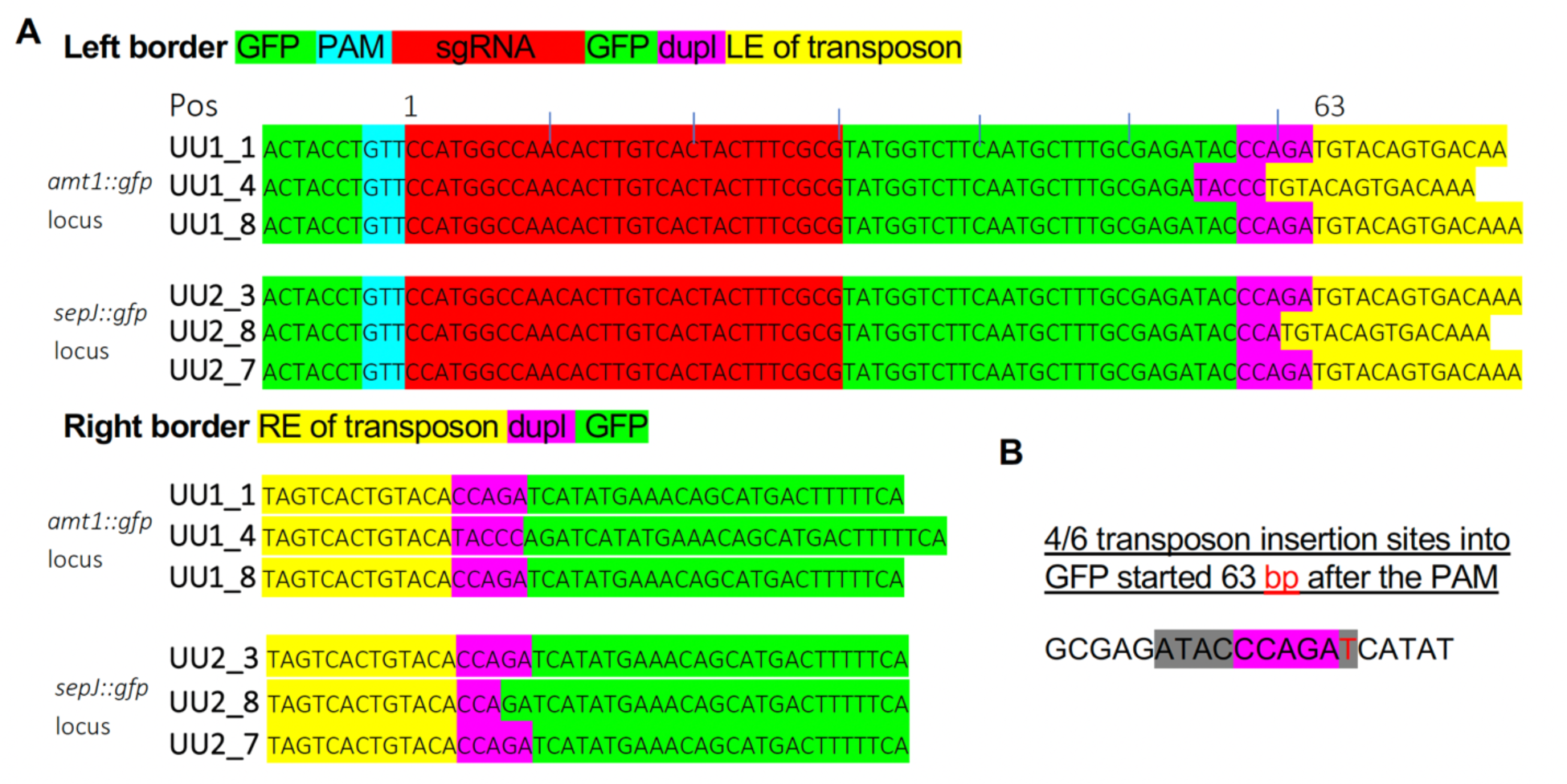
Sequences at the insertion sites of the cargo transposon after RNA-guided transposition into the *gfp* of the *amt1::gfp* or *sepJ::gfp* loci. **A,** Sequences that encode the GFP (green), or the LE and RE of the cargo transposon (yellow), are highlighted. In addition, sequences of the *gfp* that were a part of the sgRNA encoded by pAzUT.14 are highlighted that encode the PAM (blue) and the remainder sequence specific part of the sgRNA (red). Resolution of the transposase complex leads to the insertion of nucleotides, causing the small insertions highlighted in purple. The position (Pos) of the insertions was counted starting from the first base after the PAM. **B**, The insertion was exactly at position 63 for four of the six independently recovered clones derived from either parent. T in red denotes the LE start base. Similarly, resolution of the transposase led in four of the six clones to a five-base repeat, CCAGA in purple.

### RNA-guided transposition was unidirectional 5’ LE to RE 3’ and single copy insertion without co-integration of donor plasmid

To inspect the overall structure of the loci targeted by the cargo transposon in each clone sequenced, 13 kb regions of the consensus assembly sequences spanning the insertions were viewed in IGV along with aligned reads from the clone and its parent. A typical result is shown for the clone UU1.4 in Figure 6. Alignment of reads obtained from the parental strain CSVT15 revealed some single nucleotide polymorphisms between the clone and the reference strain, yet the foremost difference was the 2,918 bp insertion in the clone UU1.4 corresponding in size with the cargo transposon. Automatic annotation (Figure 6, Annotation (prokka)) identified *amt1* as *amtB* based on its homology to *amtB* from *E. coli*. Individual alignments using known sequences (Figure 6, BLAT alignments of known sequences) identified the start of the GFP, the gap caused by the insertion (line with arrows), and the remainder of *gfp*; it furthermore identified the *yfp* (VECTOR_GFPLIKE) and erythromycin resistance. Additionally, downstream of the cargo transposon insertion, it identified the vector sequences used to generate the *amt1::gfp* fusion in the parental strain CSVT15: these vector sequences inserted through a single cross-over homologous recombination event.

**Figure 6.**
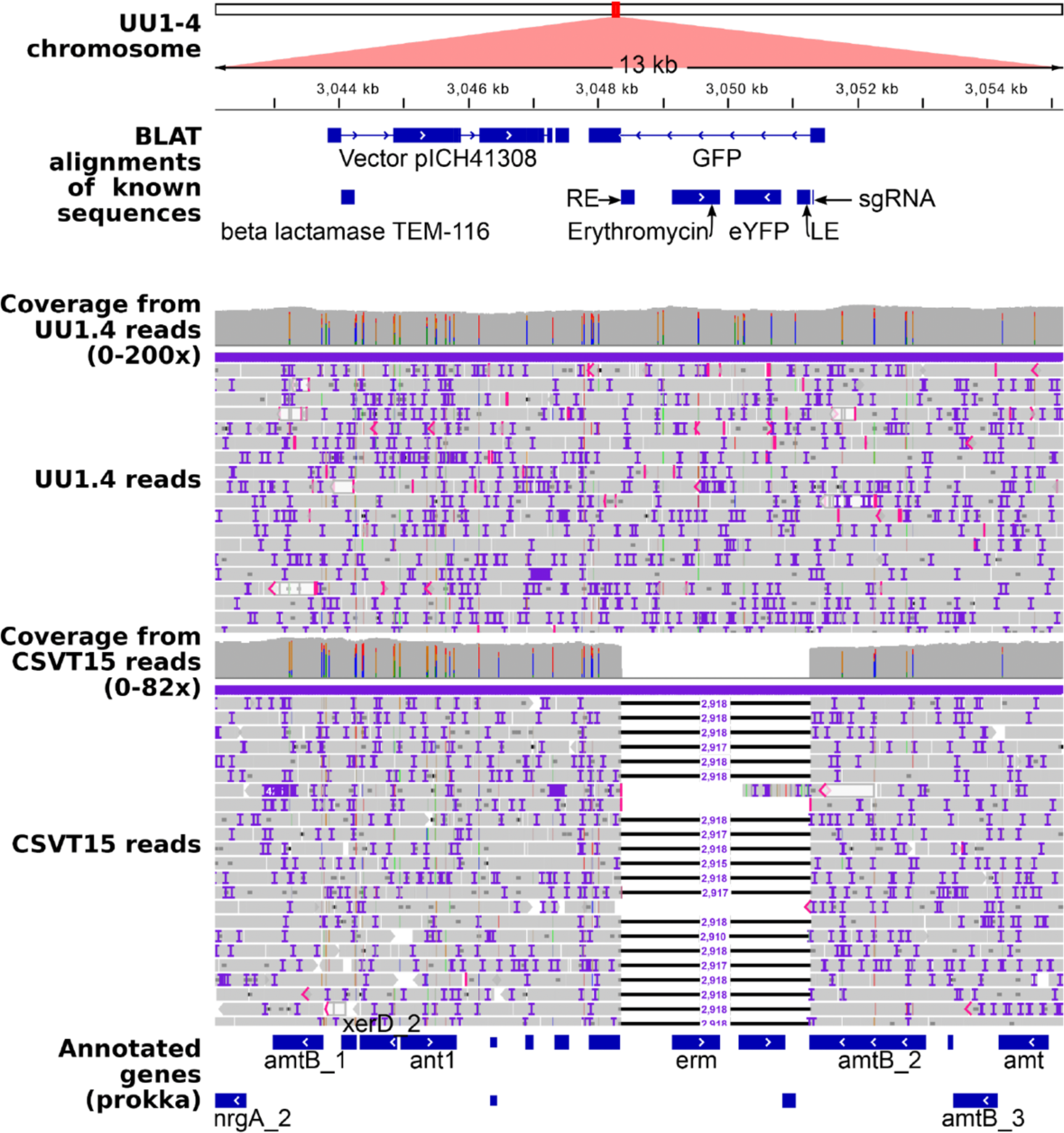
The *amt1:gfp* locus in the strain UU1.4 compared to the parental strain CSVT15. Strains were sequenced using MinIon with minimally 50 times coverage and their genomes assembled *de novo*. The assembly was automatically annotated with prokka, and known sequences were aligned using the BLAT-aligner to the *amt1::gfp* locus. In addition, MinIon reads obtained from sequencing strain UU1.4 and the parental strain CSV15 were aligned to the assembled genome of UU1.4. The assembly, automated and manual annotation, and the alignments were then visualized in Integral Genome Viewer (IGV) in the 13 kbp region spanning the 2918 bp cargo transposon and the single cross-over homologous recombination that yielded the *amt1::gfp* fusion in the parental strain CSVT15. The latter contained vector sequences 3 prime of the *gfp* (Vector pICH41308).

Results obtained from the analyses of the only six clones investigated after RNA-guided transposition revealed an identical mechanism of insertion: cargo transposon insertions guided by CAST were unidirectional from 5’LE to RE 3’ with a precise resolution of the 5’and 3’end with cut.

### No sign of off-target insertions or remobilization of endogenous mobile elements

We next examined whether exposure to the CAST machinery led to the remobilization of mobile genetic elements (MGE) already present within the genome of the parental strains. The automatic detection of indels and recombination events in genomes of the transconjugants and their respective parental strain returned, in the case of the *amt1::gfp* locus, for example, 22 putative events (Table S3). Manual inspection of the regions corresponding to these events using evidence from the aligned long-reads, however, could not verify any changes due to transposon remobilization in the nine events identified in the chromosome (Figures S6-S11). One deletion from contig_1_2444322_Sniffles2_DEL_5M4 was already present in the parental strain CSVT15 and segregated with more of less penetrance in the clones UU1.1, UU1.4 and UU1.8 (Figure S6). The events in the plasmids were mostly associated with two transposases each encoded on contig_4 and 5, respectively, the indels did not have clear borders and were difficult to evaluate as we suspected inaccuracies in the plasmid assemblies in repetitive sequences (Figure S12). We therefore conclude that exposure to the CAST machinery did not cause the remobilization of endogenous mobile genetic elements in the six cases that were analyzed in this study.

## Discussion

### CASTGATE, a Golden Gate vector set for genome engineering by RNA-guided transposition in filamentous cyanobacteria

CASTGATE uses the vocabulary from the “Golden-Gate” expanding on an existing set of vectors named CyanoGate (Vasudevan et al., 2019): we added a conjugative suicide vector, all the CAST elements and the P_glnA_ promoter for nitrogen-regulated expression restricted to cyanobacteria. In our large assemblies the expression cassette coding CAST proteins and the cargo transposon are in a single vector (Figure 1). Unlike in Rubin et al. (2021), however, we inserted the sgRNA cassette in the reverse complement orientation before the LE such that RNA polymerase on the strongly expressed sgRNA gene would not affect the transposase binding to the LE.

In addition, we designed 34 nt-long sgRNA spacers, well over the minimum required for specific targeting of RNA-guided transposition *in vitro* (Xiao et al., 2021). We included the PAM in the sgRNA spacer to test whether the binding of the sgRNA PAM sequence to the N-terminal grove of Cas12k would lead to tighter binding and therefore repression of random transposition (Xiao et al., 2021). This is possible because the PAM is not cleaved by the CAST transposase as in the case of Cas9 (Heler et al., 2015) and insertions near the PAM encoded by the sgRNA-gene would be suppressed by the LE border in the donor plasmid (Strecker et al., 2019). This configuration resulted in the specific RNA-guided integration at either of the loci targeted by the sgRNA in *Anabaena* in this study (Figures 2 and 3).

### Accurate targeting of the RNA-guided transposon cargo in *Anabaena*

In all six transconjugant *Anabaena* clones analyzed in depth, no off-target effects were detected (Table S3). These effects could have been off-target insertions of cargo DNA, or deletions and remobilization of endogenous mobile elements inside the *Anabaena* genome. Absence of off-target effects contrasted with previous reports on poor target specificity in a variety of gram-negative bacteria (Strecker et al., 2019; Rubin et al., 2021). Results from Xiao et al. (2021) showed that when Cas12 lacks the sgRNA, RNA-transposition occurs at a high rate and in random locations, therefore, the authors suggested that sgRNA binding suppresses the random transposition. In our system, sgRNA was thus expressed at a sufficiently high level such that when bound to the Cas12k it suppressed random transposon insertions at least in the small sample of analyzed clones. Higher throughput analyses will be required to investigate off-target effects in more details. The engineered CAST is toxic to the bacterial cells as *Anabaena* clones on BG11_0_ medium grew poorly when the P_glnA_ directed optimum expression of the Cas12k and an sgRNA scaffold (Figure S1).

We used the unmodified LE sequence which is suspected to direct homing via delocalized CRISPR RNA to the tRNA-leu in *S. hofmanii (sh)*: it contains a 17 bp motif matching the *sh*tRNA-Leu gene. However, in our construct the CRISPR direct repeat found upstream of the 17-bp motif in the *sh*CAST locus is missing (Saito et al., 2021). A more thorough understanding of the LE sequence is critically needed in many respects, but most importantly for the design of intron boundaries in the LE and cargo DNA so as to engineer protein fusions in coding-loci targeted by the RNA guided transposition. TnsB binding sites are the characteristic features of the LE and RE and TnsB was found to bind the backbone of the DNA only and to have a not very strict DNA-sequence preference upon binding (Kaczmarska et al., 2022).

### RNA-guided transposition was unidirectional from LE to RE, reflected a copy-and-paste mechanism, and was likely influenced by RNA

Unidirectional insertion of the cargo (Figure 5) is consistent with the unidirectional formation of the polymeric TnsC complex with the target DNA and Cas12k serving to recruit the TnsB transposase at the target site (Park et al., 2021; Querques et al., 2021).

Co-integration of the plasmid sequences supplied along with the transposon cargo was reported to occur at frequencies ranging from some 85% to 19.4% (Vo et al., 2020; Vo et al., 2021; Tou et al., 2022). It was 0.6% in the specific case of the Cas12k-homing endonuclease fusion (Tou et al., 2022). An explanation for co-integration was also proposed: the CAST lacks the TnsA required to excise the transposon and the Cas12k lacks the endonuclease activity to substitute for the TnsA. In *Anabaena*, we recovered only single copies of the cargo transposon and there were no plasmid sequence co-integrates (Figure 6). This may be due to the presence of accessory proteins in the *Anabaena* cells in which the Cas12k has evolved to be devoid of the nickase activity typically found in TnsA (Xiao et al 2021 and ref therein). Our results are of insufficient throughput to conclude definitely.

The sgRNA targeting the antisense strand of the *gfp*-gene fusions at either loci was not effective. This may have been due to the sense transcript RNA duplexing efficiently with the sgRNA and consequently less efficient RNA/DNA hetero-duplexing at the target location. For example, the gRNA depletion with DNA oligonucleotides was reported to be effective in the case of Cas9-bound gRNA (Bialk et al., 2015). The lesser efficacy of the sgRNA with a spacer containing GTT PAM may stem from a suboptimal fit at the T287 from the WED domain and the R421 from the PI domain of cas12k (Xiao et al., 2021).

### RNA-guided transposition to understand genetic features in complex communities such as symbioses

RNA-guided transposition may be used to target a locus in a specific organism in a mixture effectively: this has been recently demonstrated in bacterial communities of various complexities (Rubin et al., 2021). The advantage of this approach does not only reside in the catalysis, and therefore the efficiency with which the cargo DNA is inserted into the DNA, but also in the insertion of tags at precise locations resulting in engineered loci in cells that may be selected or followed visually in complex mixtures. We used YFP to trace whether the colonies were homogenous genetically after sonication and prolonged selection (Figure 4). The method lends itself to study fitness of the cells with engineered loci in mixed communities so as to study the role of the original versus engineered loci in microbe interactions.

We used a replicative plasmid in this initial study to be assured of sufficient expression of the CAST elements and sufficient cargo substrate for transposition. A replicative plasmid is not desired for the precise engineering of loci in microbes within complex communities because this may lead to a larger proportion of off-target insertions of the cargo. Also, curing the replicative plasmid used for the delivery of the CAST and cargo is cumbersome. We thus tested a conjugative suicide vector (pAzUT.17) for delivery of the CAST-enzymes, sgRNA and cargo DNA in a shortened protocol: after consecutive conjugation and sonication (to reduce the length of the filaments), cells were immediately transferred onto BG11_0_ medium for two days, and then on BG11-medium and selection. RNA-guided cargo DNA transposition events from conjugation with suicide plasmids were detected in clones that had no trace of the suicide plasmid (Figure S4).

We conclude that filamentous cyanobacteria, such as *Anabaena*, are amenable to genome engineering using RNA-guided transposition with the Cas12k-based CAST system. We next will need to test the approach in other species and using alternative methods of DNA-transfer. We urgently will test the method on symbiotic species, such as *Nostoc punctiforme* and *Nostoc azollae*, especially when present in complex microbial consortia such as symbioses with plants.

## Methods

### Bacterial strains and growth conditions

*Anabaena* sp. (also known as *Nostoc* sp.) strain PCC 7120 and mutants CSVT15 (Merino-Puerto et al., 2010) and CSAM137 (Flores et al., 2007) were cultured photoautrophically in BG11 or BG11_0_ (without NaNO_3_) medium (Rippka et al., 1974) at 30°C, under constant white light (35 μE×m^-2^×s^-1^) and shaking. For solid cultures, 1% (w/v) agar (Bacto-Agar, Difco) was added. When required, media were supplemented with 5 μg/ml (solid medium) or 2.5 μg/ml (liquid medium) streptomycin sulfate (Sm), spectinomycin (Sp) or erythromycin (Em).

*Escherichia coli* DH5α (Invitrogen) was used for cloning techniques, while strains HB101 and ED8634, which contain plasmid pRL623 and pRL443 (Elhai et al., 1997) respectively, were used for conjugation as described in (Elhai and Wolk, 1988). All *E. coli* strains were grown in Luria-Bertani (LB) medium supplemented with the appropriate antibiotics, incubated at 37°C and shaking for liquid cultures.

### DNA tools and vectors

*Scytonema hofmannii* Tn7-like transposase components including shCas12k, the operon encoding TnsB, TnsC and TniQ and the optimized sgRNA scaffold were obtained from pHelper_ShCAST; the left- and right-end (LE and RE) sequences of the Tn7-like transposon were obtained from pDonor_ShCAST (Strecker et al., 2019b). The pJ23119 promoter and T7Te terminator were PCR amplified from pHelper_ShCAST together with the optimized single guide (sg) RNA-scaffold sequence that contained the lgu1 sites for insertion of the target-specific spacer. The P*_glnA_* was amplified from the pRL3845 plasmid (Valladares et al., 2004). All other promoters and terminators were obtained from the CyanoGate system (Vasudevan et al., 2019). The spacer part of the sgRNA was assembled by hybridization of two complementary synthetic oligonucleotides (Integrate DNA Technologies; Figure S2 and Table S4). The coding sequences for the erythromycin and Sm/Sp resistance genes were obtained from the CyanoGate system (Vasudevan et al., 2019).

Vectors to assemble the plasmids to test CAST in cyanobacteria were obtained from the MoClo Plant Tool kit following the pipeline described in (Engler et al., 2014), where level 0 plasmids contain individual components (promoters, coding regions, terminators, etc.) and expression cassettes are assembled in level 1 plasmids. The final level T plasmids, containing all the expression cassettes, were assembled using the replicative and conjugative vector pCAT.000 or the replicative but not conjugative vector pCAT.334 form the CyanoGate system (Vasudevan et al., 2019). In addition, cyanobacterial replication encoded by OriT in pCAT.000 was replaced with the ColE1 from pEERM3 (Englund et al., 2015) to obtain a T-level backbone allowing conjugation but not replication (committing suicide) in the cyanobacterial host.

### Plasmid construct and bacterial colony screening

All plasmids generated in this work were named pAzUX.Y (plasmid Azolla Utrecht), where X indicates the level and Y is the specific ID (Table S1). They will be submitted to Addgene (reference number upon acceptance of the manuscript) to ease sharing.

To obtain plasmids pAzU0.1, pAzU0.2 pAzU0.3, sequences from the ShCAST components were PCR amplified to remove internal *Bsa*I and *Bpi*I and introduce flanking *Bsa*I restriction sites. The coding sequences for the erythromycin and Sm/Sp resistance genes were also amplified to introduce the flanking *Bsa*I sites. In all cases, Phusion High-Fidelity DNA Polymerase (ThermoFisher) was used following the manufacturer instructions. All other individual components were obtained using CyanoGate or MoClo level 0 plasmids (Figure 1). Erythromycin selection was privileged in this study because the antibiotic does not affect the plant host such as, for example, in the symbioses of ferns from the genus *Azolla* (Dijkhuizen et al., 2018).

Level 1 plasmids were assembled by digestion with *Bsa*I and ligation, following the MoClo cloning protocol (Weber et al., 2011). To obtain pAzU1.3.3, the annealed oligonucleotide spacers were introduced in plasmid pAzU1.3 containing the sgRNA scaffold by *Lgu*I digestion and ligation (Figure 1). To obtain level T plasmids, level 1 inserts were introduced in pCAT.000 or pCAT.334, together with an end-linker (L), by *Bpi*I digestion and ligation. All restriction enzymes and T4-DNA ligase were from ThermoFisher.

Plasmids were introduced in *E. coli* DH5α and HB101 by heat shock. Putative positive colonies were selected using the appropriate antibiotics. Positive colonies were confirmed by PCR with Dream Taq polymerase using specific primers. Amplified fragments were purified and sequenced (Macrogen Europe).

### Cyanobacterial transformation and sonication of exconjugants

*E. coli* strains ED8634 (containing pRL443 encoding the conjugation machinery) and HB101 (containing pRL623 and the cargo plasmid) were used for tri-parental conjugation as described in (Elhai and Wolk, 1988). The mixture of *E. coli* and cyanobacteria was spread and cultured on filters deposited on solid BG11 + 5% LB plates for 24 hours and then transferred to BG11 medium for after another 24 hours. Filters were moved to BG11 (or BG11_0_ for the rapid conjugation protocol) plates supplemented with Sm, Sp and Em for 48 hours, then transferred back to BG11 medium supplemented with the corresponding antibiotics.

Potential positive colonies growing after two weeks were re-streaked and confirmed by PCR amplification followed by sequencing, as described above.

To generate clonal strains of exconjugants, exconjugants that had integrated the YFP-encoding sequence into the genomic DNA were grown in 25 ml of BG11 medium to an optical density (OD) of 1 at 750 nm. Then, under sterile conditions, 1 ml of culture was removed and placed in a sterile plastic tube for sonication. During the sonication step, the filament-length was monitored until most of the filaments were broken down to 2-3 cells. Finally, serial dilutions were made and 50 μl of each dilution was plated on solid BG11 medium supplemented with the appropriate antibiotic and allowed to grow under the conditions described above. The genotypes of the clonal colonies thus obtained were analyzed by PCR (Table S4).

To test relative growth rates, cultures grown in liquid BG11 medium with the corresponding antibiotic for 1 week were washed with BG11_0_ medium, inoculated with the chlorophyll equivalents indicated and incubated in the light at 30°C for 8, 12 and 28 days.

### Confocal microscopy

Samples were grown on a plate of BG11 medium supplemented with the corresponding antibiotic under conditions previously described. Biomass was taken with a toothpick and suspended in 100 μl of sterile distilled water. For fluorescence detection, drops of 10 μl were placed on a new BG11 plate, cut out of the agar and covered with a cover slip. Images were photographed using a Leica SP5 microscope (40X oil immersion objective). The YFP was excited using a 514 nm laser and the sf-GFP using a 488 nm laser, both lasers were used at 20% power and the irradiation came from an argon ion laser. The fluorescence of YFP was visualized with a window of 515-545 nm, for sf-GFP a window of 500-525 nm was used. Autofluorescence from the natural pigments of cyanobacterial cells was collected using a window of 640-740 nm. ImageJ software was then used to remove background as well as overlapping images (Schindelin et al., 2015).

### DNA extraction and sequencing

Cyanobacterial genomic DNA was isolated using the GeneJET Genomic DNA Purification Kit (ThermoFisher) following the manufacturer’s instructions. DNA quality and concentration were analyzed by UV absorption, q-bit and on gel. Minion sequencing libraries were generated using 0.3-3 µg DNA with the SQK-LSK109 kit following instructions by Nanopore Technologies (version NBE_9065_v109_revAK_14Aug2019); in case of multiplexing the EXP-NBD104 extension was combined with NEB Blunt/TA Ligase Master Mix (M0367, New England BioLabs (NEB)). Briefly, the DNA was first repaired (NEBNextFFPE Repair mix (M6630) and NEBNext Ultra II End repair/dA-tailing Module (E7546)), then bound on AMPure XP beads (Beckman & Coulter) and cleaned twice with 75% v/v ethanol, before elution in water at 50°C for 10 min. Subsequently, when barcoded, the barcode adapters were ligated and the DNA cleaned once more using the AMPure XP beads and 75% v/v ethanol. Finally, the DNA was ligated to the sequencing primers in the presence of the tether, and washed with Short Fragment Buffer (SQK-LSK109 kit, Nanopore Technologies) bound once more on the AMPure XP beads, before elution in elution buffer at 50°C for 10 min. Priming and loading of the recycled MinION flowcell (R9.4.1 Nanopore Technologies, starting with 600 active pores) was as per the manufactuers instruction using reagents from EXP-FLP002 (Nanopore Technologies). The flowcell was washed between the loading of different libraries using the EXP-WSH004 (Nanopore Technologies) reagents that included DNase.

### Genome assemblies and annotation

Data acquisition of the nanopore sequencing used the Minknow program (Oxford Nanopore Technologies) until 50 times genome coverage was achieved for the barcode with the lowest reads. The actual coverage ranged from 51 (UU1_1) to 233 (UU1_4), with two outliers 21 (CSAM) and 35 (UU2_3). Base calling was carried out separately. Assemblies were computed *de novo* using Flye (Kolmogorov et al., 2019) with default settings, then visualized using Bandage (Wick et al., 2021); polishing using medaka proved not to improve the assemblies; annotation of the assemblies was carried out with bakta (Schwengers et al., 2021) and prokka (Seemann, 2014) for comparison. *Anabaena* genomes, and alignment files of the minion reads aligned to them with minimap2 (Li, 2018), were visualized using Integral Genome Viewer (IGV; Thorvaldsdóttir et al., 2013). The BLAT function inside the IGV was used to locate YFP, GFP, Amt1, SepJ as well as the left and right end sequences of the Tn7 provided in the original plasmid pAzUT14. Large structural variations between the transconjugant after RNA-guided transposition and the reference genomes were programmatically detected using Sniffles 2, after mapping of the reads using NGMLR (Sedlazeck et al., 2018). The log of the analyses is detailed at https://github.com/lauralwd/anabaena_nanopore_workflow/blob/main/script.sh. Sniffles 2 is highly dependent on assembly quality and we therefore show results for assemblies with the highest coverage. When comparing all of the strains with the transposon targeted into the *amt1::gfp* locus, Sniffles 2 identified 15 insertions, 3 deletions and 4 recombinations (Table S3); 13 putative events were in the plasmids. Given the higher fluidity with which plasmids were assembled, we suspected over-sampling during assembly, but read coverage for the plasmids was similar to that of the chromosome. The indels were then inspected by extracting the FASTA files defined by their boundaries in Sniffles 2, then locating their regions in the assemblies of the transconjugants using BLAT in IGV for further evaluation (Figures 5 to 12).

### Material and Data availability

CASTGATE vectors listed in Table S1 have been deposited with Addgene (reference numbers provided upon manuscript acceptance). Minion sequencing data were deposited at the European Nucleotide Archive (ENA) and are made available under the accession number PRJEB60371.

### Accession numbers

The DNA sequencing data from the article may be found at the ENA accession number PRJEB60371.

## Supporting information

The following supporting information is available for publication with figures documenting, the relative toxicity of the CASTGATE plasmids in Anabaena

## Acknowledgements

We thank Gracia Benítez for maintaining the cyanobacterial stocks and cultures. We further thank the Gordon and Betty Moore Foundation’s Symbiosis in Aquatic Systems Initiative (Prime Contract No. 9355) for funding, the Netherlands Science Organization (NWO-ALWGS.2016.020) for funding L.W. Dijkhuizen and University of Alcalá for supporting DA financially at Utrecht University.

## Author contributions

SN-B, PL and HS conceived and oversaw the project. EF provided reference strains and material. SA, DPR, DA carried out the cloning, SA generated the transconjugants and SA and CS-B verified RNA-guided transposition by PCR and confocal microscopy. EF analyzed the growth fitness of the exconjugants. HS and LWD sequenced, assembled and analyzed the genomes. All authors contributed to generating and writing the manuscript and agreed to its final form.

## Supporting Information for publication

The following supporting information is available for publication with figures documenting, the relative toxicity of the CASTGATE plasmids in *Anabaena*, the sgRNA that target *gfp,* and results from conjugation with a CASTGATE suicide plasmid. In addition, figures are provided to evaluate candidate alterations in the genomes from the strains obtained after exposure to the CAST enzymes. Furthermore, tables are provided that identify the CASTGATE vectors cloned and those tested, a list of indels computationally identified in genomes from strains obtained after exposure to CAST enzymes, and the list of primers used in this study.

**Figure S1** Relative toxicity of the CASTGATE plasmids in *Anabaena*.

**Figure S2** The three different sgRNAs targeting *gfp*.

**Figure S3** Efficient targeting at locus *alr3727* in wild-type *Anabaena*.

**Figure S4** Rapid conjugation protocol for RNA-guided transposition using the suicide plasmid pAzUT.17.

**Figure S5** Visualization in IGV of the chromosome loci for Indels 1 (contig_1_86_Sniffles2_INS_0M4) and 2 (contig_1_4114_Sniffles2_INS_1M4).

**Figure S6** Visualization in IGV of the locus for indel 3 (contig_1_2350996_Sniffels2.DEL_4M4).

**Figure S7** Visualization in IGV of the locus for indel 4 (contig_1_2444322_Sniffles2_DEL_5M4).

**Figure S8** Visualization in IGV of the locus for indel 5 (contig_1_4325042_Sniffles2_INS_6M4).

**Figure S9** Visualization in IGV of the locus for indel 6 (contig_1_4886060_Sniffles2_BND_8M4).

**Figure S10** Visualization in IGV of the *amt1::gfp* locus for indels 7 (contig_1_5088816_Sniffles2_DEL_AM4) and 8 (contig_1_5089223_Sniffles2_INS_9M4).

**Figure S11** Visualization in IGV of the locus for indel 9 (contig_1_6211681_Sniffles2_INS_BM4).

**Figure S12** Visualization in IGV of the locus for indel 10 (contig_3_26226_Sniffles2_INS_0M1), typical of the indels detected by Snifles 2 in the plasmid sequences.

**Table S1** CASTGATE vectors generated in this study.

**Table S2** CASTGATE vectors transferred to and tested in *Anabaena* wild-type, and the CSVT15 and CSAM137 strains in this study.

**Table S3** Insertions detected by “Snifles2” comparing genome assemblies from the parental strains with those from clones obtained after RNA-guided transposition.

**Table S4** Primers used for PCR assays and key cloning steps in this study.

## References

1. Baldanta S, Guevara G, Navarro-Llorens JM (2022) SEVA-Cpf1, a CRISPR-Cas12a vector for genome editing in cyanobacteria. Microbial Cell Factories. 21:1–3.

2. Bialk P, Rivera-Torres N, Strouse B, Kmiec EB (2015) Regulation of gene editing activity directed by single-stranded oligonucleotides and CRISPR/Cas9 systems. PLoS One 10: e0129308

3. Dijkhuizen LW, Brouwer P, Bolhuis H, Reichart GJ, Koppers N, Huettel B, Bolger AM, Li FW, Cheng S, Liu X, Wong GK, Pryer K, Weber A, Bräutigam A, Schluepmann H (2018) Is there foul play in the leaf pocket? The metagenome of floating fern *Azolla* reveals endophytes that do not fix N_2_ but may denitrify.” New Phytol 217: 453–466.

4. Elhai J, Vepritskiy A, Muro-Pastor AM, Flores E, Wolk CP (1997) Reduction of conjugal transfer efficiency by three restriction activities of *Anabaena* sp. strain PCC 7120. J Bacteriol 179: 1998–2005

5. Elhai J, Wolk CP (1988) [83] Conjugal transfer of DNA to cyanobacteria. Methods Enzymol 167: 747–754

6. Engler C, Youles M, Gruetzner R, Ehnert TM, Werner S, Jones JDG, Patron NJ, Marillonnet S (2014) A Golden Gate modular cloning toolbox for plants. ACS Synth Biol 3: 839–843

7. Englund E, Andersen-Ranberg J, Miao R, Hamberger B, Lindberg P (2015) Metabolic engineering of *Synechocystis* sp. PCC 6803 for production of the plant diterpenoid manoyl oxide. ACS Synth Biol 4: 1270–1278

8. Faure G, Shmakov SA, Yan WX, Cheng DR, Scott DA, Peters JE, Makarova KS, Koonin E v. (2019) CRISPR–Cas in mobile genetic elements: counter-defence and beyond. Nature Reviews Microbiology 2019 17:8 17: 513–525

9. Flores E, Pernil R, Muro-Pastor AM, Mariscal V, Maldener I, Lechno-Yossef S, Fan Q, Wolk CP, Herrero A (2007) Septum-localized protein required for filament integrity and diazotrophy in the heterocyst-forming cyanobacterium *Anabaena* sp. strain PCC 7120. J Bacteriol 189: 3884–3890

10. Gutiérrez S, Lauersen KJ, Fabris M, Abbriano R (2021) Gene delivery technologies with applications in microalgal genetic engineering. Biology 2021, Vol 10, Page 265 10: 265

11. Heler R, Samai P, Modell JW, Weiner C, Goldberg GW, Bikard D, Marraffini LA (2015) Cas9 specifies functional viral targets during CRISPR–Cas adaptation. Nature 2015 519:7542 519: 199–202

12. Hoffmann FT, Kim M, Beh LY, Wang J, Vo PLH, Gelsinger DR, George JT, Acree C, Mohabir JT, Fernández IS, et al (2022) Selective TnsC recruitment enhances the fidelity of RNA-guided transposition. Nature 2022 1–10

13. Hu B, Yang G, Zhao W, Zhang Y, Zhao J (2007) MreB is important for cell shape but not for chromosome segregation of the filamentous cyanobacterium *Anabaena* sp. PCC 7120. Mol Microbiol 63: 1640–1652

14. Jiang F (2017) CRISPR-Cas9 Structures and Mechanisms. Article in Annual Review of Biophysics. doi: 10.1146/annurev-biophys-062215-010822

15. Kaczmarska Z, Czarnocki-Cieciura M, Górecka-Minakowska KM, Wingo RJ, Jackiewicz J, Zajko W, Poznański JT, Rawski M, Grant T, Peters JE, et al (2022) Structural basis of transposon end recognition explains central features of Tn7 transposition systems. Mol Cell 82: 2618–2632.e7

16. Klompe SE, Vo PLH, Halpin-Healy TS, Sternberg SH (2019) Transposon-encoded CRISPR–Cas systems direct RNA-guided DNA integration. Nature 571: 219–225

17. Kolmogorov M, Yuan J, Lin Y, Pevzner PA (2019) Assembly of long, error-prone reads using repeat graphs. Nature Biotechnology 2019 37:5 37: 540–546

18. Li H (2018) Minimap2: pairwise alignment for nucleotide sequences. Bioinformatics 34: 3094–3100

19. Merino-Puerto V, Mariscal V, Mullineaux CW, Herrero A, Flores E (2010) Fra proteins influencing filament integrity, diazotrophy and localization of septal protein SepJ in the heterocyst-forming cyanobacterium Anabaena sp. Mol Microbiol 75: 1159–1170

20. Mutalipassi M, Riccio G, Mazzella V, Galasso C, Somma E, Chiarore A, De Pascale D, Zupo V (2021) Symbioses of cyanobacteria in marine environments: Ecological insights and biotechnological perspectives. mdpi.com. doi: 10.3390/md19040227

21. Niu TC, Lin GM, Xie LR, Wang ZQ, Xing WY, Zhang JY, Zhang CC (2018) Expanding the potential of CRISPR-Cpf1-based genome editing technology in the cyanobacterium Anabaena PCC 7120. ACS Synth Biol 8:170–80.

22. Park JU, Tsai AWL, Mehrotra E, Petassi MT, Hsieh SC, Ke A, Peters JE, Kellogg EH (2021) Structural basis for target site selection in RNA-guided DNA transposition systems. Science 373: 768–774.

23. Peters J, development NC-G&, 2001 undefined (2001) Tn7 recognizes transposition target structures associated with DNA replication using the DNA-binding protein TnsE. genesdev.cshlp.org. doi: 10.1101/gad.870201

24. Peters JE, Craig NL (2001) Tn7: smarter than we thought. Nature Reviews Molecular Cell Biology 2001 2:11 2: 806–814

25. Querques I, Schmitz M, Oberli S, Chanez C, Jinek M (2021) Target site selection and remodelling by type V CRISPR-transposon systems. Nature 2021 599:7885 599: 497–502

26. Rikkinen J (2017) Cyanobacteria in terrestrial symbiotic systems. Modern Topics in the Phototrophic Prokaryotes: Environmental and Applied Aspects 243–294

27. Rippka R, Waterbury J, Cohen-Bazire G (1974) A cyanobacterium which lacks thylakoids. Archives of Microbiology 1974 100:1 100: 419–436

28. Rodrigo González Linares B, Rita Costa Cristóbal Almendros Romero Stan J Brouns Thesis committee AJ, J Brouns Chirlmin Joo Christophe Danelon SJ (2020) A CRISPR-associated transposase presents null cargo integration efficiency when targeting a transcriptionally highly active region.

29. Rubin BE, Diamond S, Cress BF, Crits-Christoph A, Lou YC, Borges AL, Shivram H, He C, Xu M, Zhou Z, et al (2021) Species- and site-specific genome editing in complex bacterial communities. Nature Microbiology 2021 7:1 7: 34–47

30. Saito M, Ladha A, Strecker J, Faure G, Neumann E, Altae-Tran H, Macrae RK, Zhang F (2021) Dual modes of CRISPR-associated transposon homing. Cell 184: 2441–2453.e18

31. Sánchez-Baracaldo P, Bianchini G, Wilson JD, Knoll AH (2022) Cyanobacteria and biogeochemical cycles through Earth history. Trends in Microbiology 30: 143– 157

32. Schindelin J, Rueden CT, Hiner MC, Eliceiri KW (2015) The ImageJ ecosystem: An open platform for biomedical image analysis. Mol Reprod Dev 82: 518–529

33. Schwengers O, Jelonek L, Dieckmann MA, Beyvers S, Blom J, Goesmann A (2021) Bakta: rapid and standardized annotation of bacterial genomes via alignment-free sequence identification. Microb Genom. doi: 10.1099/MGEN.0.000685

34. Sedlazeck FJ, Rescheneder P, Smolka M, Fang H, Nattestad M, von Haeseler A, Schatz MC (2018) Accurate detection of complex structural variations using single-molecule sequencing. Nat Methods 15: 461–468

35. Seemann T (2014) Prokka: rapid prokaryotic genome annotation. Bioinformatics 30: 2068–2069

36. Stellwagen AE, Craig NL (1997) Avoiding self: Two Tn7-encoded proteins mediate target immunity in Tn7 transposition. EMBO Journal 16: 6823–6834

37. Strecker J, Ladha A, Gardner Z, Schmid-Burgk JL, Makarova KS, Koonin E v., Zhang F (2019) RNA-guided DNA insertion with CRISPR-associated transposases. Science (1979). doi: 10.1126/science.aax9181

38. Thorvaldsdóttir H, Robinson JT, Mesirov JP (2013) Integrative Genomics Viewer (IGV): high-performance genomics data visualization and exploration. Brief Bioinform 14: 178–192

39. Tou CJ, Orr B, Kleinstiver BP (2022) Cut-and-Paste DNA Insertion with Engineered Type V-K CRISPR-associated Transposases. bioRxiv 2022.01.07.475005

40. Ungerer J, Pakrasi HB (2016) Cpf1 is a versatile tool for CRISPR genome editing across diverse species of cyanobacteria. Sci rep 6:39681.

41. Valladares A, Muro-Pastor AM, Herrero A, Flores E (2004) The NtcA-dependent P1 promoter is utilized for glnA expression in N2-fixing heterocysts of *Anabaena* sp. strain PCC 7120. J Bacteriol 186: 7337–7343

42. Vasudevan R, Gale GAR, Schiavon AA, Puzorjov A, Malin J, Gillespie MD, Vavitsas K, Zulkower V, Wang B, Howe CJ, et al (2019) CyanoGate: A Modular Cloning Suite for Engineering Cyanobacteria Based on the Plant MoClo Syntax. Plant Physiol 180: 39–55

43. Vo PLH, Acree C, Smith ML, Sternberg SH (2021) Unbiased profiling of CRISPR RNA-guided transposition products by long-read sequencing. Mob DNA. doi: 10.1186/S13100-021-00242-2

44. Vo PLH, Ronda C, Klompe SE, Chen EE, Acree C, Wang HH, Sternberg SH (2020) CRISPR RNA-guided integrases for high-efficiency, multiplexed bacterial genome engineering. Nature Biotechnology 2020 39:4 39: 480–489

45. Weber E, Engler C, Gruetzner R, Werner S, Marillonnet S (2011) A Modular Cloning System for Standardized Assembly of Multigene Constructs. PLoS One 6: e16765

46. Wick R, Schultz M, Zobel J, Bioinformatics KH-, 2015 undefined Bandage: interactive visualization of de novo genome assemblies. academic.oup.com

47. Xiao R, Wang S, Han R, Li Z, Gabel C, Mukherjee IA, Chang L (2021) Structural basis of target DNA recognition by CRISPR-Cas12k for RNA-guided DNA transposition. Mol Cell 81: 4457–4466.e5

48. Zehr JP, Capone DG (2020) Changing perspectives in marine nitrogen fixation. Science. doi: 10.1126/SCIENCE.AAY9514

49. Zhang S, Guo F, Yan W, Dai Z, Dong W, Zhou J, Zhang W, Xin F, Jiang M (2020) Recent Advances of CRISPR/Cas9-Based Genetic Engineering and Transcriptional Regulation in Industrial Biology. Frontiers in Bioengineering and Biotechnology 7: 459

