## Supplementary material for "Genome engineering by RNA-guided transposition for *Anabaena* PCC 7120": The following supporting information is available for publication with figures documenting, the relative toxicity of the CASTGATE plasmids in Anabaena

**Title:**

**Affiliations:**

**Current affiliation DA:**

[5] Department of Life Sciences, University of Alcalá, Alcalá de Henares, Spain.

The following supporting information is available for publication with figures documenting, the relative toxicity of the CASTGATE plasmids in *Anabaena*, the sgRNA that target *gfp,* and results from conjugation with a CASTGATE suicide plasmid. In addition, figures are provided to evaluate candidate alterations in the genomes from the strains obtained after exposure to the CAST enzymes. Furthermore, tables are provided that identify the CASTGATE vectors cloned and those tested, a list of indels computationally identified in genomes from strains obtained after exposure to CAST enzymes, and the list of primers used in this study.

**Figure S1** Relative toxicity of the CASTGATE plasmids in *Anabaena*.

**Figure S2** The three different sgRNAs targeting *gfp*.

**Figure S3** Efficient targeting at locus *alr3727* in wild-type *Anabaena*.

**Figure S4** Rapid conjugation protocol for RNA-guided transposition using the suicide plasmid pAzUT.17.

**Figure S5** Visualization in IGV of the chromosome loci for Indels 1 (contig_1_86_Sniffles2_INS_0M4) and 2 (contig_1_4114_Sniffles2_INS_1M4).

**Figure S6** Visualization in IGV of the locus for indel 3 (contig_1_2350996_Sniffels2.DEL_4M4).

**Figure S7** Visualization in IGV of the locus for indel 4 (contig_1_2444322_Sniffles2_DEL_5M4).

**Figure S8** Visualization in IGV of the locus for indel 5 (contig_1_4325042_Sniffles2_INS_6M4).

**Figure S9** Visualization in IGV of the locus for indel 6 (contig_1_4886060_Sniffles2_BND_8M4).

**Figure S10** Visualization in IGV of the *amt1::gfp* locus for indels 7 (contig_1_5088816_Sniffles2_DEL_AM4) and 8 (contig_1_5089223_Sniffles2_INS_9M4).

**Figure S11** Visualization in IGV of the locus for indel 9 (contig_1_6211681_Sniffles2_INS_BM4).

**Figure S12** Visualization in IGV of the locus for indel 10 (contig_3_26226_Sniffles2_INS_0M1), typical of the indels detected by Snifles 2 in the plasmid sequences.

**Table S1** CASTGATE vectors generated in this study.

**Table S2** CASTGATE vectors transferred to and tested in *Anabaena* wild-type, and the CSVT15 and CSAM137 strains in this study.
**Table S3** Insertions detected by “Snifles2” comparing genome assemblies from the parental strains with those from clones obtained after RNA-guided transposition.
**Table S4** Primers used for PCR assays and key cloning steps in this study.

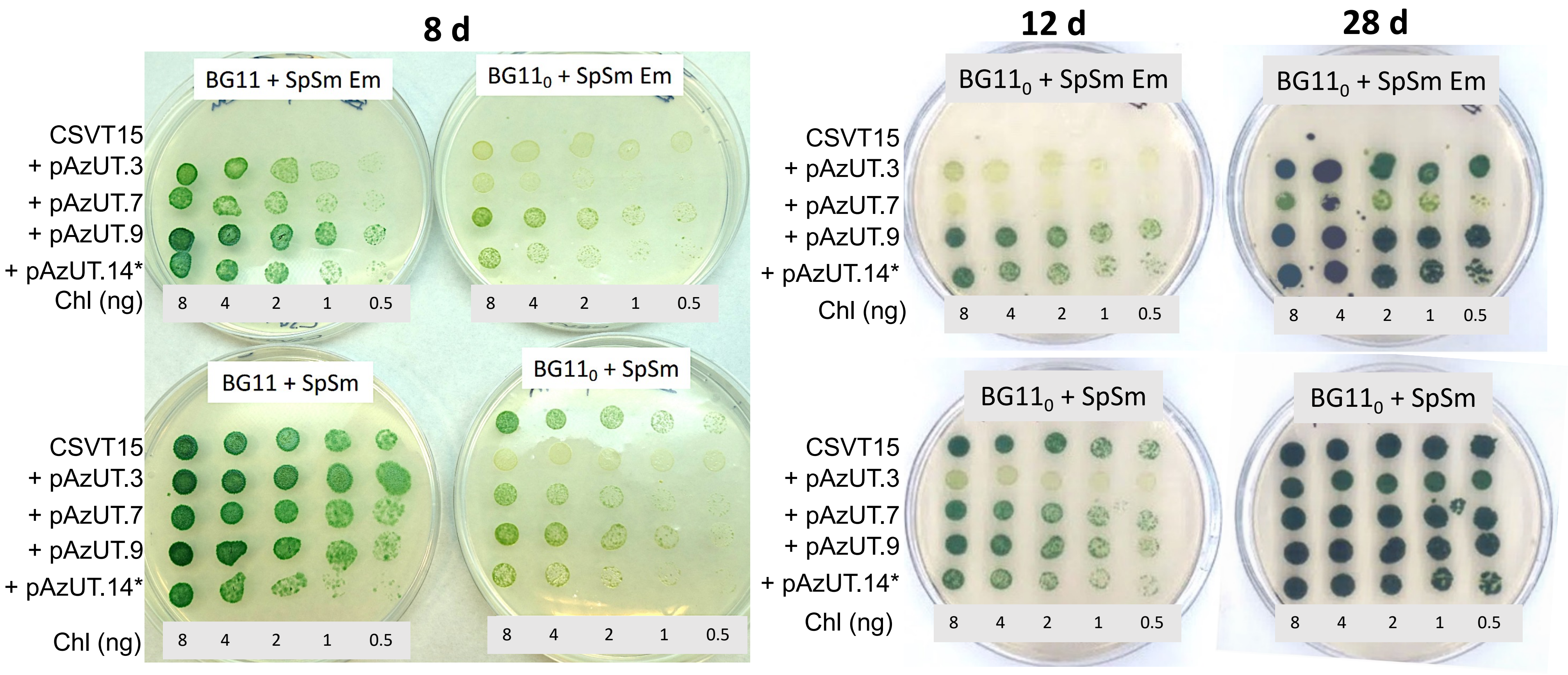


**Figure S1 Relative toxicity of the CAST plasmids in *Anabaena*.** Exconjugant strains derived from CSVT15 generated by conjugation contained the following constructs: pAzUT.3, with only the YFP and erythromycin-resistance cassettes expressed constitutively; pAzUT.7 with, in addition to the components of pAzUT.3, the $P_{glnA}$-driven *cas12k* expression induced under nitrogen deprivation, and the strongly and constitutively expressed sgRNA scaffold (without the target specific sequence); pAzUT.9, with $P_{glnA}$-driven expression of all the CAST genes (*tnsB*, *tnsC*, *tniQ*, and *cas12k*), the cargo transposon flanked by the LE and RE, and the strongly and constitutively expressed sgRNA-scaffold without a target sequence. The strain carrying pAzUT.14*, used as a control, was homozygous for the cargo transposon inserted into the *gfp* and was cured of the pAzUT.14 plasmid that contained, in addition to what is encoded by pAzUT.9, the sgRNA targeting sense sequence of *gfp*. Strains with the plasmids and the controls, the parental strain and pAzUT.14 were grown in successive dilutions of starting material, measured in chlorophyll *a* equivalents (Chl in ng), for 8, 12 and 28 days on either Sm/Sp or Sm/Sp and Em antibiotic selection in BG11 medium (with nitrogen) and BG11$_0$ medium (without combined nitrogen).

A

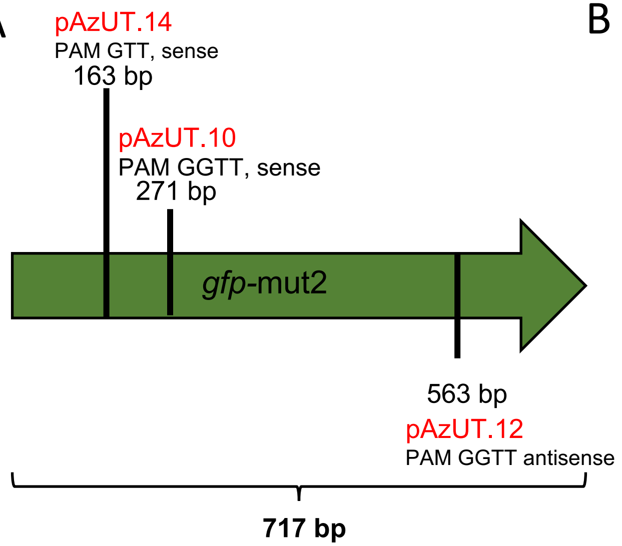


B

| p label | Primer Orientation | Sequence | PAM | Target Strand |
|---|---|---|---|---|
| pAzUT.10 | Fwd | TT**GCTCTTC**CAAA**GGTT**ATGTACAGGAAAGAACTATATTTTTCAAAGGCTG**GAAGAGC**AA | **GGTT** | Sense |
| | Rev | TT**GCTCTTC**CAGCCTTTGAAAAATATAGTTCTTTCCTGTACATAACCTTTG**GAAGAGC**AA | | |
| pAzUT.12 | Fwd | TT**GCTCTTC**CAAA**GGTT**GTCTGGTAAAAGGACAGGGCCATCGCCAATGCTG**GAAGAGC**AA | **GGTT** | Antisense |
| | Rev | TT**GCTCTTC**CAGCATTGGCGATGGCCCTGTCCTTTTACCAGACAACCTTTG**GAAGAGC**AA | | |
| pAzUT.14 | Fwd | TT**GCTCTTC**CAAA**GTT**CCATGGCCAACACTTGTCACTACTTTCGCGGCTG**GAAGAGC**AA | **GTT** | Sense |
| | Rev | TT**GCTCTTC**CAGCCGCGAAAGTAGTGACAAGTGTTGGCCATGGAACTTTG**GAAGAGC**AA | | |

**Figure S2 The three different sgRNAs targeting *gfp*.** A, Schematic representation of the coding region of *gfp-mut2* showing the base at which the different target-specific sequences start that are encoded by the sgRNA. For example, the target specific sequence of the sgRNA encoded in pAzUT.14 starts 163 bp inside the 717 bp-long *gfp-mut2* sequence with the GTT PAM sequence and encodes the sense strand. B, Primers used for cloning of the target specific sequences into the LguI restriction site of the scaffold sgRNA from pAzU1.3 (depicted in Fig.1). Forward (Fwd) and reverse (Rev) primers were annealed, then in one pot with pAzU1.3, digested and ligated to obtain the target specific sgRNA from the pAzUT.10, 12 and 14 as indicated. LguI restriction site (red), orientation specific overhangs (green), PAM (blue).

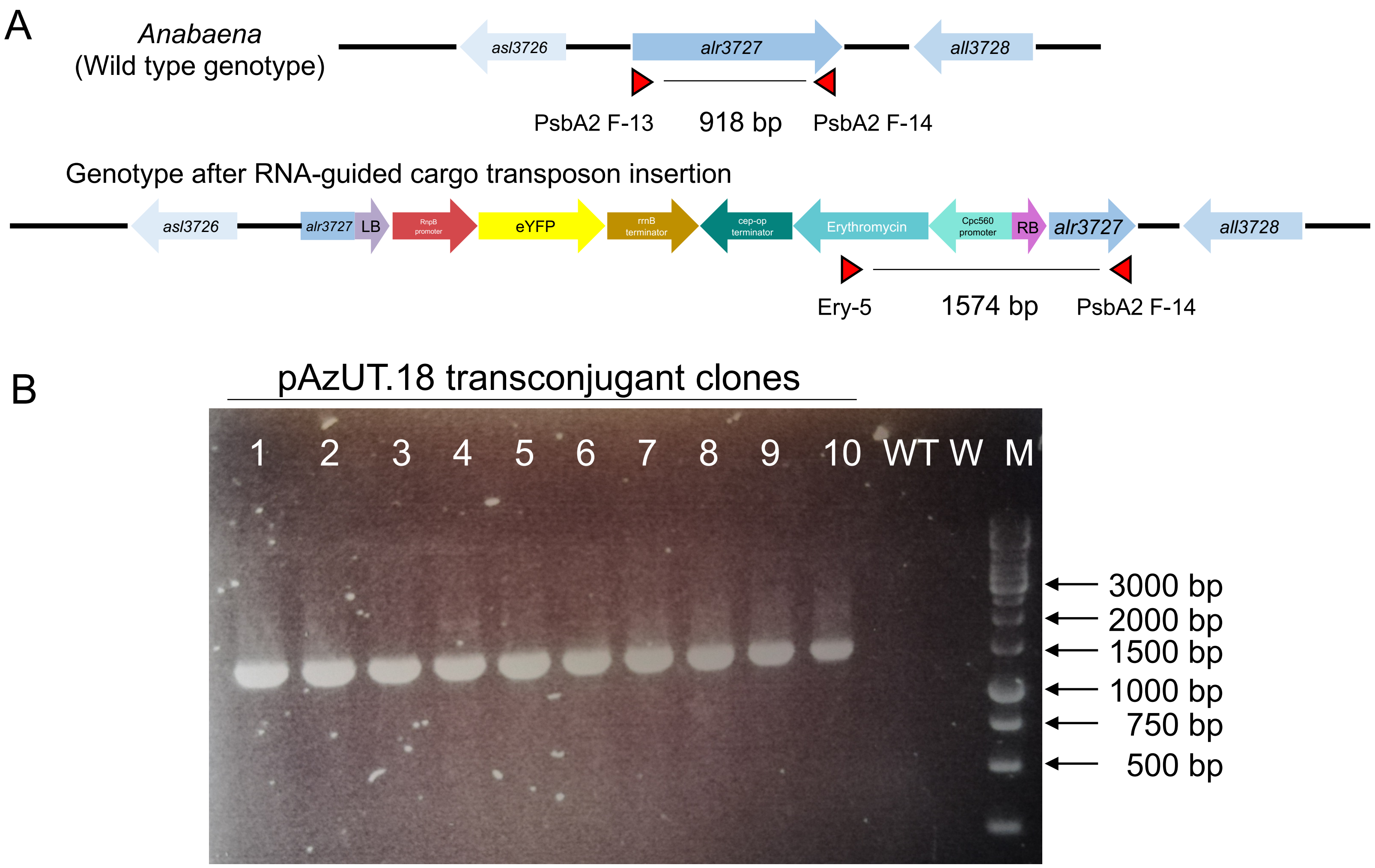


**Figure S3 Efficient targeting at locus *alr3727* in wild-type *Anabaena*.** A, PCR strategy to detect RNA-guided transposition of the cargo transposon encoded in pAzUT.18 into the locus alr3727 of *Anabaena* sp. PCC 7120. (B) Fragments amplified from transconjugant clones obtained using a rapid process as follows: conjugation, subsequent transfers for 48 h on $BG11_0$ medium, followed by sonication and then selection on $BG11_0$ medium with erythromycin. 1-10, randomly selected transconjugant clones; WT, *Anabaena* sp. PCC 7120; W, water (negative control); M, DNA molecular weight standards.

A

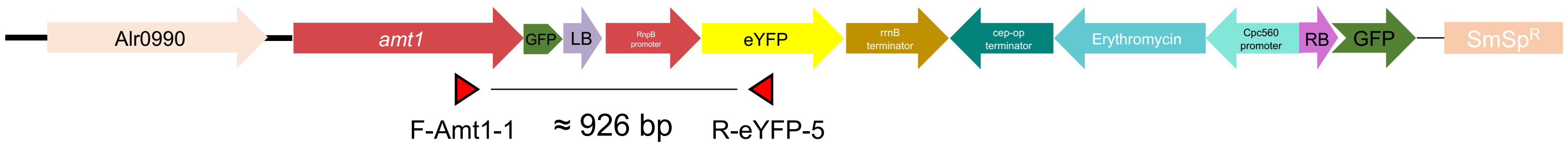


B

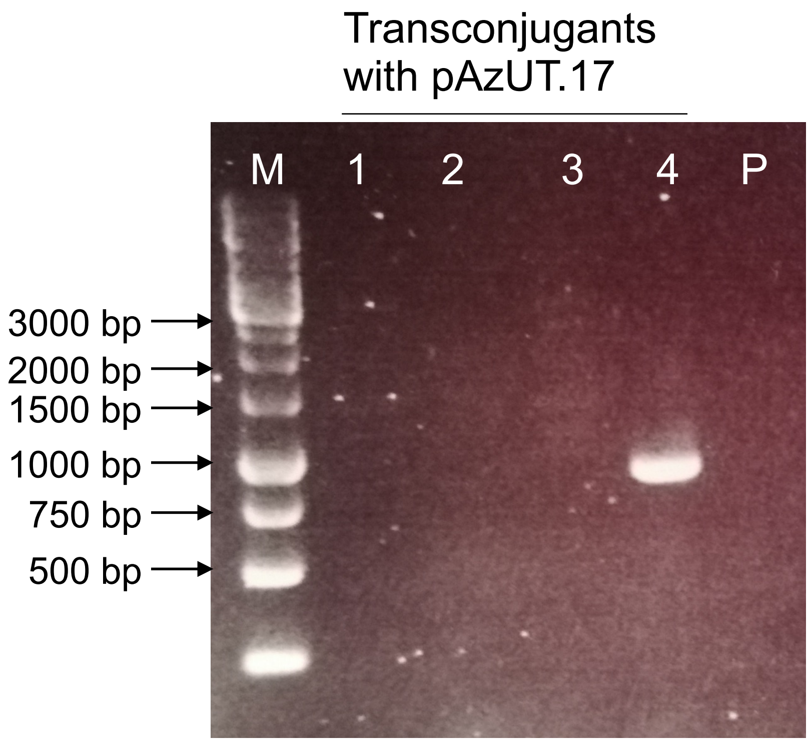


**Figure S4 Rapid conjugation protocol for RNA-guided transposition using the suicide plasmid pAzUT.17**. pAzUT.17 is identical to pAzUT.14 except in the plasmid backbone region where the cyanobacterial replication elements were removed. In the rapid conjugation protocol with the parental strain CSV15, material spread in a filter on $BG11_0$ medium was transferred to $BG11_0$ medium during the first 48 h after of conjugation. Thereafter, transconjugants were selected on BG11 medium supplemented with erythromycin. A, Scheme of the *gfp* after insertion of RNA-guided cargo transposon and the PCR used to detect the insertion. B, PCR detection of the insertion at the LE of the cargo transposon. M, DNA molecular weight markers; 1-4, transconjugants with pAzUT.17 derived from CSV15 ; P, parental strain, CSVT15 (*amt1::gfp*).

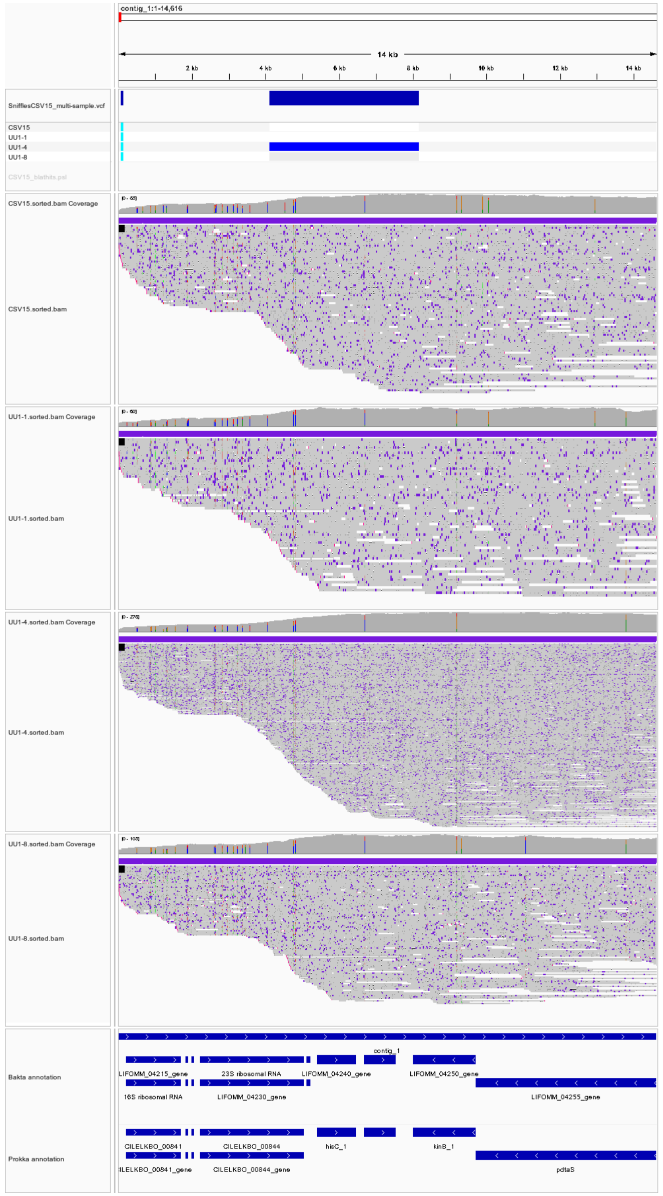


**Figure S5 Visualization in IGV of the chromosome loci for Indels 1 (contig_1_86_Sniffles2_INS_0M4) and 2 (contig_1_4114_Sniffles2_INS_1M4).** These insertions are at the border of the chromosome and have no distinct border, therefore, they are a likely result of assembly issues at the end of the chromosome.

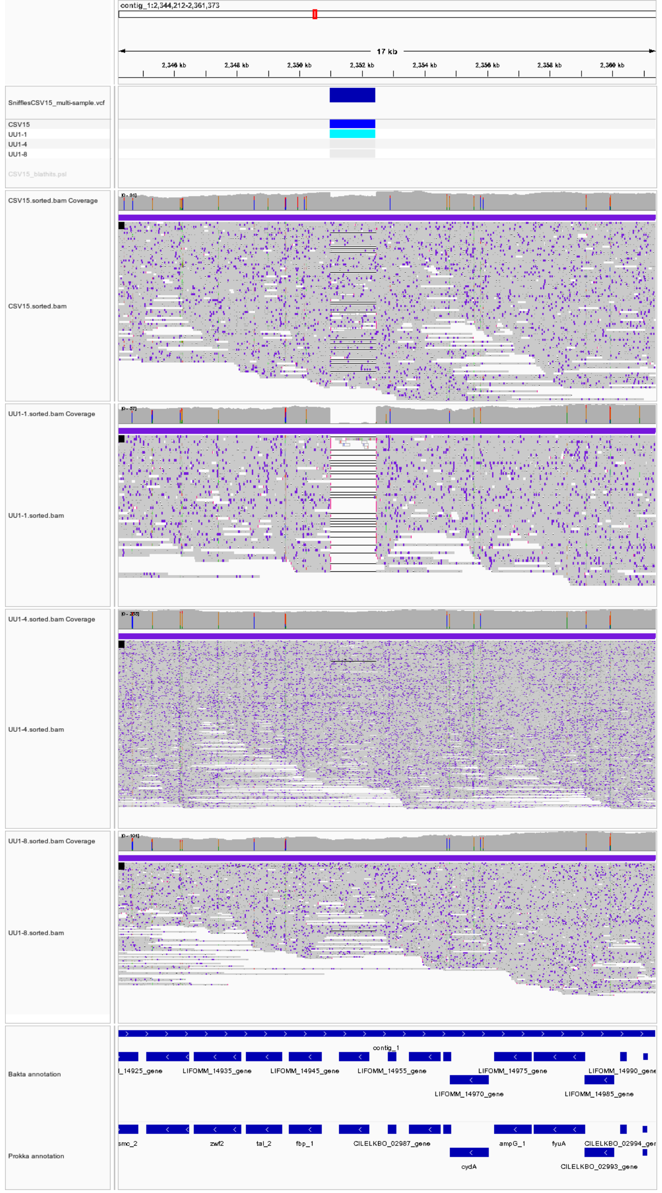


**Figure S6 Visualization in IGV of the locus for indel 3 (contig_1_2350996_Sniffels2.DEL_4M4).** The deletion is already partially found in the parental strain CSVT15; it is dominant in the UU1_1 but barely detectable in the UU1_4 or UU1_8. This indel suggests relative instability of the chromosome of *Anabaena*, consistent with our finding when comparing the genome assemblies from CSVT15 and wt (data not shown).

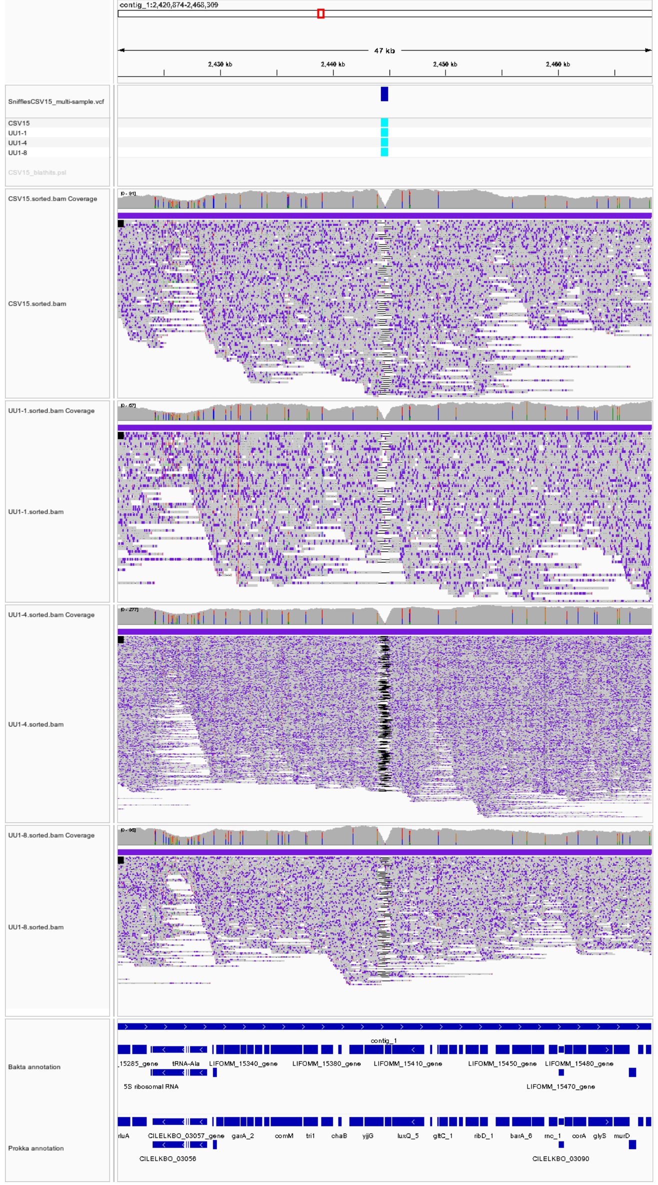


**Figure S7 Visualization in IGV of the locus for indel 4 (contig_1_2444322_Sniffles2_DEL_5M4).** The v-shape lack of coverage for this deletion in all strains sequenced suggests a problem with assembly of the chromosome in the parental strain, possibly due to a region difficult to sequence. False positive indel.

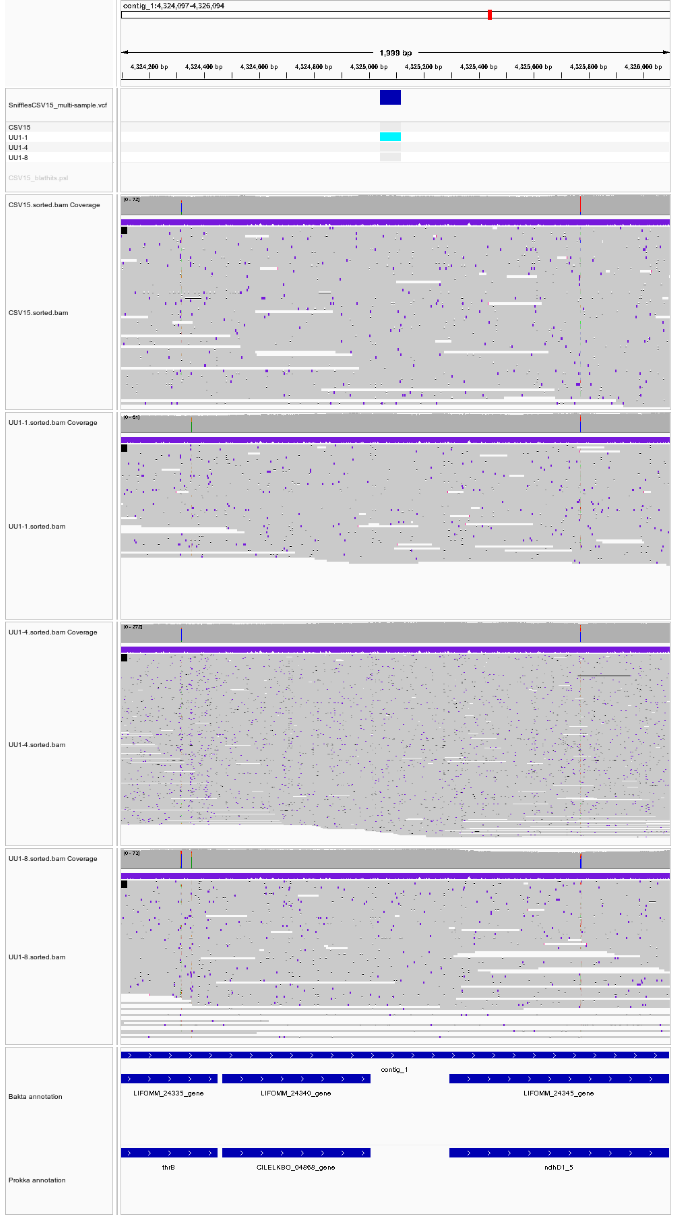


**Figure S8 Visualization in IGV of the locus for indel 5 (contig_1_4325042_Sniffles2_INS_6M4).** Only a very minor proportion of the reads support the recombination in this region with differing end points, false positive indel thus.

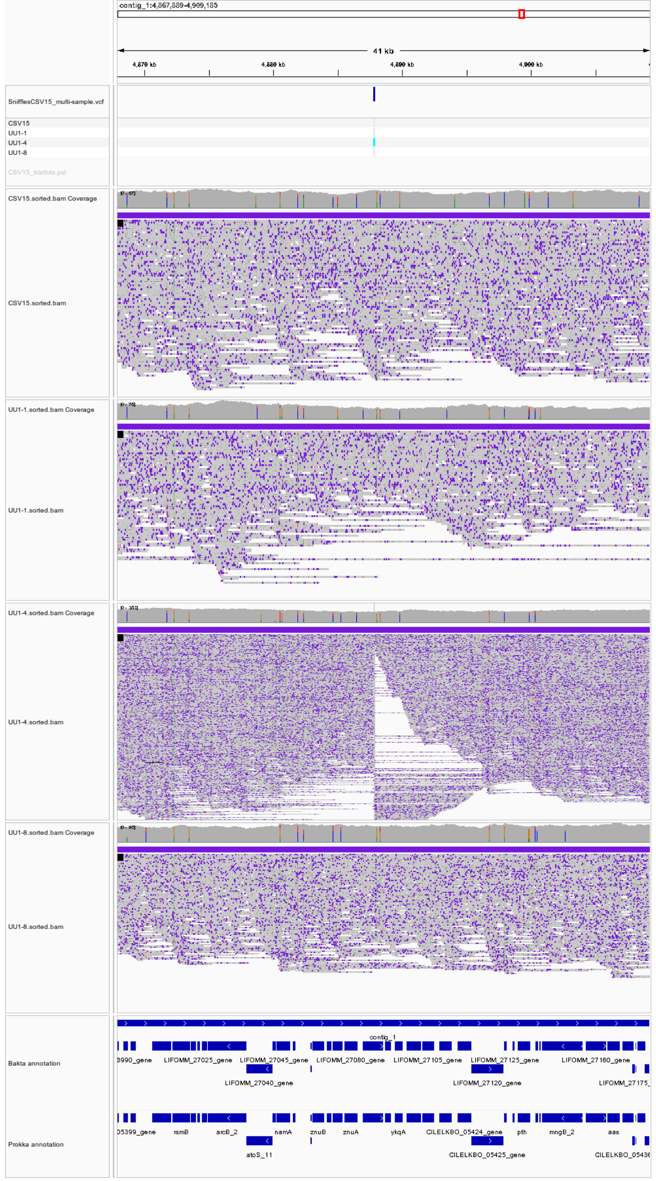


**Figure S9 Visualization in IGV of the locus for indel 6 (contig_1_4886060_Sniffles2_BND_8M4).**

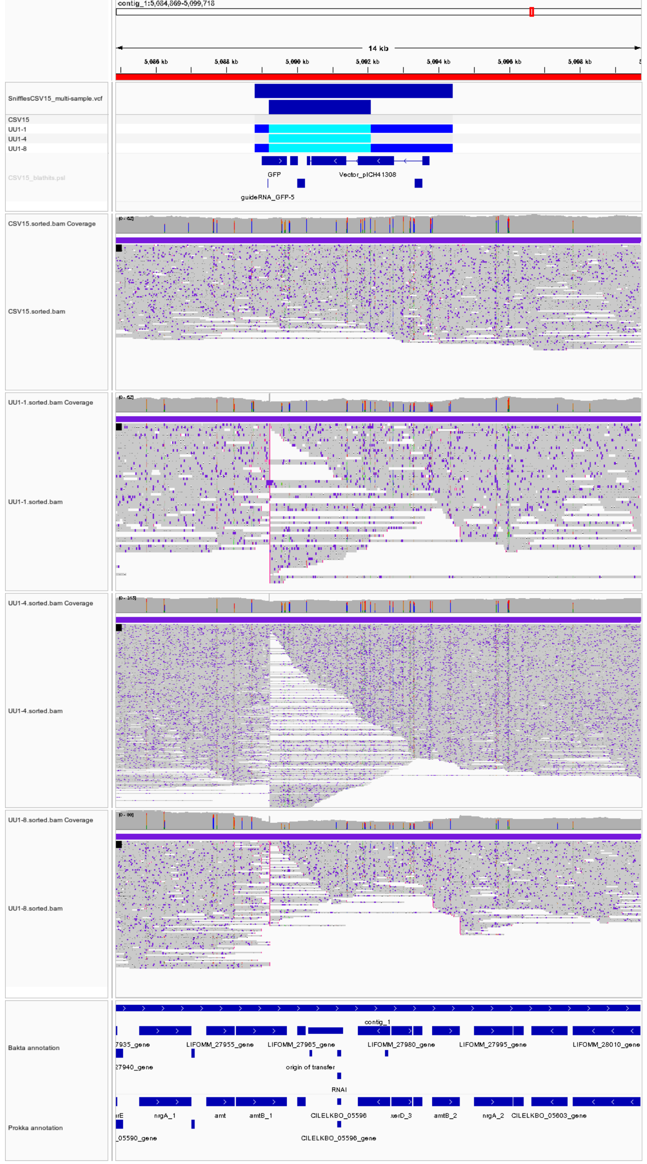


**Figure S10 Visualization in IGV of the *amt1::gfp* locus for indels 7 (contig_1_5088816_Sniffles2_DEL_AM4) and 8 (contig_1_5089223_Sniffles2_INS_9M4).** In UU1-1 and UU1-8, reads were cut sharply at several locations of the plasmid sequences that were inserted when CSVT15 was generated by single cross-over recombination, these are the deletions for indel 7. Penetrance of these events were weak, however. The alignment of the reads is cut sharply and the location of the cargo transposon insertion inside the GFP sequence with 100% penetrance in UU1-1,4 and 8 but not in the parental, this is the true positive insertion from indel 8.

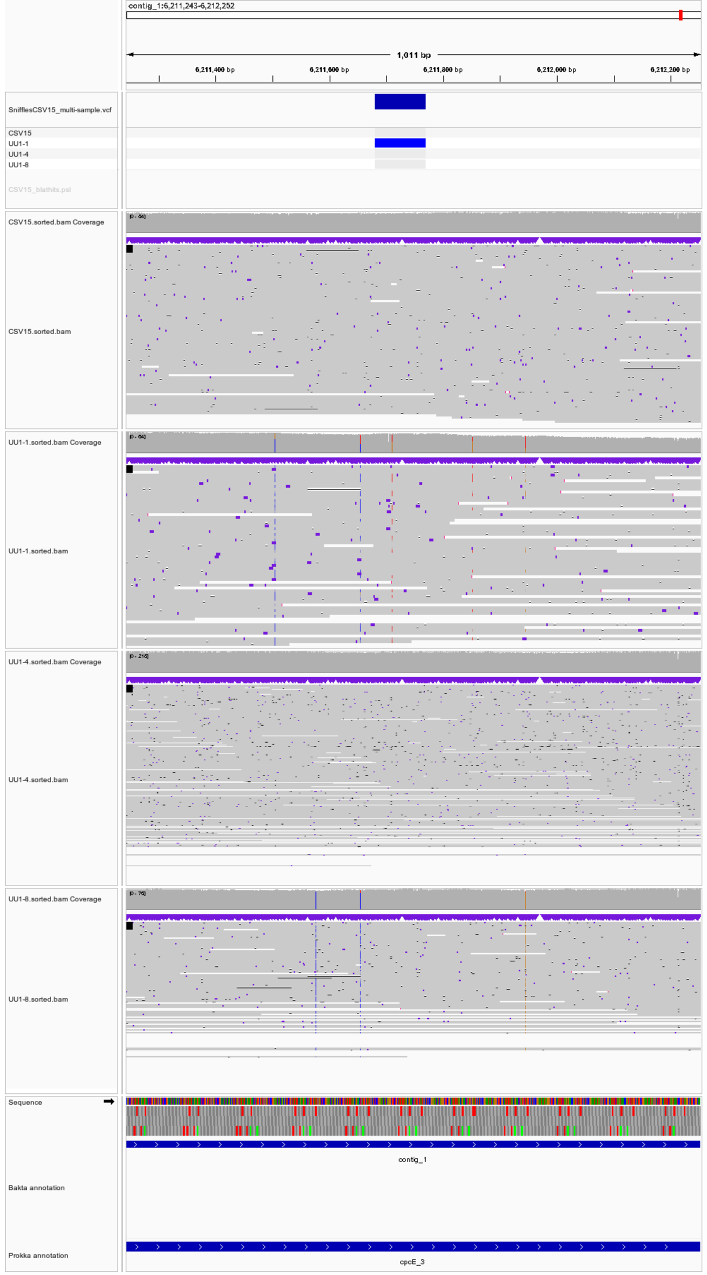

**Figure S11 Visualization in IGV of the locus for indel 9 (contig_1_6211681_Sniffles2_INS_BM4).** The insertion is short and not supported by many reads.

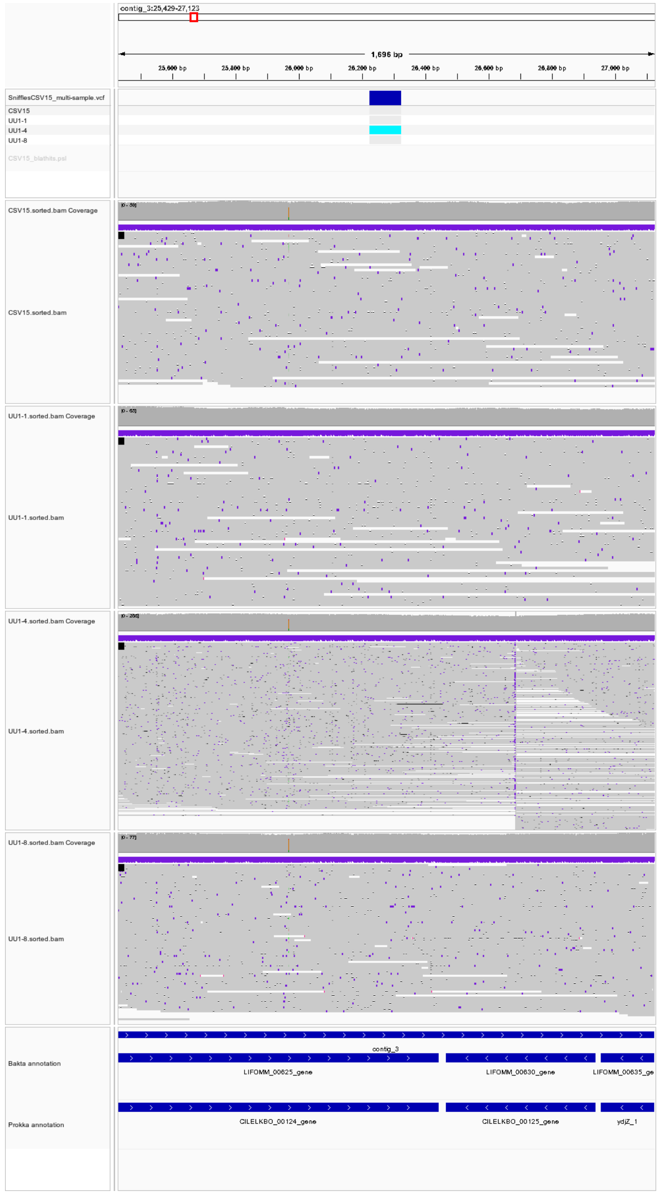


**Figure S12 Visualization in IGV of the locus for indel 10 (contig_3_26226_Sniffles2_INS_0M1), typical of the indels detected by Snifles 2 in the plasmid sequences**. The indels in plasmids more often than not were located in repetitive sequences encoding transposase.

| | Strains | | |
|---|---|---|---|
| Plasmid (brief description) | Anabaena PCC7120 (WT) | CSVT15 (amt1::gfp) | CSAM137 (sepJ::gfp) |
| pAzUT.3 (only eYFP and AB cassette) Erythromycin R | ✓ | ✓ | ✓ |
| pAzUT.4 (only eYFP and AB cassette) Sp/Sm R | ✓ | ✓ | ✓ |
| pAzUT.5 (- Target, LB and RB) | ✓ | ✓ | ✓ |
| pAzUT.6 (- Target, LB and RB) | ✓ | | |
| pAzUT.7 (- Tns and target) | ✓ | ✓ | ✓ |
| pAzUT.8 (- Tns and target) | ✓ | | |
| pAzUT.9 (- Target) | ✓ | ✓ | ✓ |
| pAzUT.10 (gGFP option 1) | | ✓ | ✓ |
| pAzUT.12 (gGFP option 2) | ✓ | ✓ | ✓ |
| pAzUT.14 (gGFP option 3) | ✓ | ✓ | ✓ |
| pAzUT.17 (Conjugative suicide and gGFP 3) | | ✓ | ✓ |
| pAzUT.18 (pCAT.000 gPsbA2) | ✓ | | |
| pAzUT.19 (pCAT.000 gUrtA) | ✓ | | |
| pAzUT.20 (Conjugative suicide and gGFP 1) | | ✓ | ✓ |

**Table S2** CASTGATE vectors transferred to and tested in *Anabaena* wild-type, and the CSVT15 and CSAM137 strains in this study. CSVT15 had been generated by single cross-over recombination so as to generate the *amt1::gfp* fusion (Merino-Puerto et al., 2010) and grew on $BG11_0$ medium without nitrogen. Similarly, CSAM137 had been generated by recombination so as to generate the *sepJ::gfp* fusion (Flores et al., 2007); CSAM137 grew only slowly if at al on the $BG11_0$ medium.

| Insertion nr | type | genomic region | sequence ID | length | CSVT15 | UU1.1 | UU1.4 | UU1.8 | IGV quality control | annotation |
|---|---|---|---|---|---|---|---|---|---|---|
| 1 | insertion | chromosome | contig_1_86_Sniffles2_INS_0M4 | 172 | X | X | X | X | noise at contig extremity | transposase |
| 2 | insertion | chromosome | contig_1_4114_Sniffles2_INS_1M4 | 8189 | | | X | | noise at contig extremity | Trichormus hypothetical protein |
| 3 | deletion | chromosome | contig_1_2350996_Sniffels2.DEL_4M4 | -1467 | X | X | | | present already in CSVT15 | IS110 transposase |
| 4 | deletion | chromosome | contig_1_2444322_Sniffles2_DEL_5M4 | -693 | X | X | X | X | assembly issue in CSVT15 reference | Histidine kinase response regulator |
| 5 | insertion | chromosome | contig_1_4325042_Sniffles2_INS_6M4 | 77 | | X | | | false positive | no good hit |
| 6 | recombination | chromosome | contig_1_4886060_Sniffles2_BND_8M4 | NA | | | X | | true in UU1.4 | NA |
| 7 | deletion | chromosome | contig_1_5088816_Sniffles2_DEL_AM4 | -5613 | | X | | X | true in UU1.1 and UU1.8 | homologous recombination of GFP plasmid |
| 8 | insertion | chromosome | contig_1_5089223_Sniffles2_INS_9M4 | 2893 | | X | X | X | positive | CAST insertion |
| 9 | insertion | chromosome | contig_1_6211681_Sniffles2_INS_BM4 | 89 | | X | | | false positive | HEAT repeat domain containing protein / glycosyltransferase |
| 10 | insertion | plasmid | contig_3_26226_Sniffles2_INS_0M1 | 99 | | | X | | false positive | IS5 transposase |
| 11 | insertion | plasmid | contig_4_56839_Sniffles2_INS_0M0 | 1658 | | X | | | positive | ISNCY transposase |
| 12 | insertion | plasmid | contig_4_82848_Sniffles2_INS_1M0 | 905 | | X | | | positive | no good hit |
| 13 | insertion | plasmid | contig_5_49595_Sniffles2_INS_0M2 | 205 | | X | | | positive | - |
| 14 | insertion | plasmid | contig_5_139017_Sniffles2_INS_2M2 | 1672 | | X | | | positive | NIES-23 transposase |
| 15 | recombination | plasmid | contig_5_148784_Sniffels2_BND_5M2 | NA | | | X | | positive | transposase |
| 16 | recombination | plasmid | contig_5_151176_Sniffels2_BND_5M2 | NA | | | X | | positive | same transposase as above |
| 17 | insertion | plasmid | contig_5_174398_Sniffles2_INS_3M2 | 221 | | | X | | positive | ISNCY transposase |
| 18 | insertion | plasmid | contig_5_193393_Sniffles2_INS_4M2 | 1666 | | | | X | positive | |
| 19 | insertion | plasmid | contig_5_260548_Sniffles2_INS_7M2 | 1671 | | X | | | positive | ISNCY transposase |
| 20 | insertion | plasmid | contig_5_266694_Sniffles2_INS_8M2 | 1467 | | X | | | positive | ISNCY transposase |
| 21 | recombination | plasmid | contig_5_267538_Sniffles2_BND_9M2 | NA | | | | X | positive | |
| 22 | insertion | plasmid | contig_5_399380_Sniffles2_INS_AM2 | 1670 | | | X | | positive | ISNCY transposase |

**Table S3 All the indels detected by Snifles 2 comparing genome assemblies from the parental strain with those from clones obtained after RNA-guided transposition in the *amt1::gfp* locus**. Sequence identity was specified starting with the contig in which it was encoded. Contig_1 encoded the chromosome. X identifies detection of the indel in either the parental (CSVT15) or the strains with the RNA-guided insertion of the cargo transposon in the *amt1::gfp* locus (UU1.1, UU1.4 and UU1.8). Contig_2- to contig_5 encompassed the plasmids (plasmid assemblies seemed more fluid, possibly because of assembly issues). The analyses were highly dependent on the quality of the assemblies, hence the quality control in IGV to visualize the sequence reads aligned to the parental CSVT15 genome assembly (IGV quality control). We provide the IGV-visualizations to substantiate conclusions for all the 9 indels detected in the chromosome in Supplemental Figures S5-S11, we also provide the example for a plasmid indel 10 in Supplemental Figure S12.

| Name | Sequence | PCR-assay/cloning step |
|---|---|---|
| **Amt1-1** | GAGTCACCAGAGAAGAAGAATTGGA | Fig.2 PCR1 and PCR3 |
| **Ery-5** | CCTGATGAATGAGGGTAACACG | Fig. 1 and Fig.2 PCR2 |
| **eYFP-5** | GCAGATGAACTTCAGGGTCAG | Fig.1 and Fig.2 PCR1 |
| **GFP-3** | CCATGCCATGTGTAATCCCAG | Fig.1 and Fig. 2 PCR2 |
| **SepJ-1** | CCCAAATATTCGGAGTTTTATTCGTCA | Fig.1 PCR1 and PCR3 |
| **pCAT.000-10** | GATACCTTGTGCGGCTATGT | Fig.1 and Fig.2 PCR4 |
| **pCAT.000-11** | GTTTGGTTGATGCGAGTGATTT | |
| **Ery1-F** | TTGAAGACAAAATGAATAAAAATATTAAATACTCTC | Amplification of erythromycin resistance gene CDS from pC0.029 |
| **Ery2-R** | TTGAAGACAAAAGCTTATTTCCGCCCATTAAACAAC | |
| **ProGlnA_1** | TTGAAGACTTGGAGCGCATTCCTTCTTCTC | Amplification of the $P_{GlnA}$ for insertion into Level 0 Promoter + 5 UTR |
| **ProGlnA_2** | TTGAAGACAAATGGTGTTACTCCTTCTCTGCCA | Amplification of the $P_{GlnA}$ for insertion into Level 0 Promoter + 5 UTR |
| **ProGlnA_3** | TTGAAGACAACATTTGTTACTCCTTCTCTGCCA | Pair with ProGlnA_1 to clone Pro+5U with ATG |
| **Spe1-F** | TTGAAGACAAAATGCGCGAAGCGGTTATTG | Amplification of spectinomycin resistance gene CDS from pC0.028 |
| **Spe2-R** | TTGAAGACAAAAGCTTACTTGCCGACCACTTTTG | |
| **LB-Bsa-F1** | TTGGTCTCAAGCGAGGCGTAGTGTACAGTGAC | Amplification of Left End from pDONOR (Strecker et al., 2019) for the Level 1 construct |
| **LB-Bsa-R2** | TTGGTCTCAGGAGTCAGTAACTATTAAACTTAGGGGTGGG | |
| **LB-Bsa-F3** | TTGGTCTCAGGAGAGGCGTAGTGTACAGTG | Amplification of Left End from pDONOR (Strecker et al., 2019) to fuse to the Promoter-eYFP-Terminator cassette |
| **LB-Bsa-R4** | TTGGTCTCAAGTATCAGTAACTATTAAACTTAGGGGTG | |
| **RB-Bsa-F1** | TTGGTCTCAAGCGAAGGCGACAGTCAATTTGTC | Amplification of Right End from pDONOR (Strecker et al., 2019) for the Level 1 construct |
| **RB-Bsa-R2** | TTGGTCTCAGGAGCTACGTCTCTACGTGTACAG | |
| **RB-Bsa-F4** | TTGGTCTCATACTAAGGCGACAGTCAATTTGTC | Amplification of Left End from pDONOR (Strecker et al., 2019) to fuse to a Promoter-EryR-Terminator cassette. Use with RB-Bsa-R2 |
| **gGFP-1 (F1)** | TTGCTCTTCCAAAGGTTATGTACAGGAAAGAACTATATTTTTCAAAGGCTGGAAGAGCAA | gRNA of GFP-Mut2 using GGTT as PAM sequence and restriction enzyme LguI. R1 is a complement strand of F1. Used for pAzUT.10 |
| **gGFP-2 (R1)** | TTGCTCTTCCAGCCTTTGAAAAATATAGTTCTTTCCTGTACATAACCTTTGGAAGAGCAA | |
| **gGFP-3 (F2)** | TTGCTCTTCCAAAGGTTGTCTGGTAAAAGGACAGGGCCATCGCCAATGCTGGAAGAGCAA | gRNA targeting GFP-Mut2 using GGTT as PAM sequence in complement strand, and restriction enzyme LguI. R2 is a complement strand of F2. Used for pAzUT.12 |
| **gGFP-4 (R2)** | TTGCTCTTCCAGCATTGGCGATGGCCCTGTCCTTTTACCAGACAACCTTTGGAAGAGCAA | |
| **gGFP-5 (F3)** | TTGCTCTTCCAAAGTTCCATGGCCAACACTTGTCACTACTTTCGCGGCTGGAAGAGCAA | gRNA of GFP-Mut2 using GTT as PAM sequence and restriction enzyme LguI. R3 is a complement strand of F3. Used for pAzUT.14 |
| **gGFP-6 (R3)** | TTGCTCTTCCAGCCGCGAAAGTAGTGACAAGTGTTGGCCATGGAACTTTGGAAGAGCAA | |

**Table S4 Primers used for PCR assays and key cloning steps in this study.**
